# Pitch motor areas contribute to the perception of prosodic categories in speech

**DOI:** 10.64898/2026.06.22.733802

**Authors:** Seung-Cheol Baek, Seung-Goo Kim, Burkhard Maess, Daniela Sammler

## Abstract

Prosody is a fundamental aspect of speech characterized by suprasegmental features such as pitch. Prosodic pitch contours convey speakers’ intentions, for example, by distinguishing statements from questions. Understanding these intentions requires abstracting continuous, variable pitch information into discrete categories. The perception of segmental speech features has been proposed to recruit motor areas according to the articulatory effectors used to produce them. However, whether this principle extends to vocal pitch and prosodic category perception remains unclear. To address this question, we collected magnetoencephalography data from 29 participants (15 females) while they identified the prosody (Statement vs. Question) of single words varying in pitch contour along a five-level continuum; 25 participants also completed a singing localizer involving simple five-tone melodies. Region-restricted searchlight decoding of singing versus rest identified two premotor regions relevant to vocal pitch production, anatomically corresponding to the putative ventral and dorsal laryngeal motor cortex. Perceived prosodic categories were decodable from these regions, with a stronger contribution from the dorsal subregion more closely linked to pitch modulation, particularly in the left hemisphere. Moreover, decoding performance mirrored behavioral discriminability across the prosody continuum before typical response times, but not later, suggesting functional relevance to perceptual categorization. Finally, bidirectional transfer of category-related information between pitch motor and auditory areas argued against these motor areas merely echoing processing in auditory regions. Together, these findings support effector-relevant motor area involvement in prosodic category perception, thereby broadening the scope of motor contributions to speech perception from vocal-tract articulators to the laryngeal sound source.

**Significance Statement:** Speech perception has been proposed to recruit the premotor cortex, with subregions linking speech sounds to the articulatory gestures that produce them. However, whether this principle extends to prosody—the pitch changes distinguishing statements from questions—remains unclear. Using magnetoencephalography, we localized premotor regions relevant to pitch production during singing and examined them during prosody perception. Neural patterns in these regions distinguished perceived prosodic categories and tracked listeners’ categorization of clear and ambiguous prosody before typical response times, but not later, suggesting behavioral relevance. These regions also exchanged category-related information with auditory areas, indicating that their activity did not merely echo processing in auditory regions. Our findings extend motor area involvement from articulators to the sound source underlying prosodic pitch.

## Introduction

The role of the premotor cortex (PMC) in speech perception has long been debated. Classical motor theories posit that perception relies critically on mapping speech sounds onto articulatory gestures in an effector-specific manner (Liberman and Mattingly, 1985). For example, lip and tongue motor regions may contribute to identifying labial and dental sounds, respectively, thereby providing perceptual references for categorical speech units (e.g., /b/ and /d/; D’Ausilio et al., 2009). In contrast, more recent models suggest that speech perception primarily relies on the ventral auditory stream (e.g., Hickok and Poeppel, 2007; DeWitt and Rauschecker, 2012), and that the PMC merely provides supplementary support for sensory analysis or auditory-motor integration under specific listening or task conditions, although evidence for such involvement remains mixed (e.g., Hickok, 2010; Du et al., 2014; Perron et al., 2024). However, this debate has largely centered on segmental features shaped by supralaryngeal articulators, which configure the vocal-tract filter, leaving open whether motor area involvement extends to suprasegmental features generated through control of the laryngeal sound source (Fant, 1960).

Prosody, characterized by suprasegmental features including pitch and loudness, is another fundamental aspect of speech crucial for verbal communication (Pierrehumbert and Hirschberg, 1990). Particularly, prosodic pitch contours signal speakers’ intentions (Hellbernd and Sammler, 2016; Kurumada and Roettger, 2022), which listeners decode by abstracting continuous, variable acoustic information into discrete categories—for instance, distinguishing questions from statements (Xie et al., 2021). Vocal pitch is produced by laryngeal muscle movements, which are dually represented in ventral and dorsal subregions of the PMC, collectively termed the laryngeal motor cortex (LMC; e.g., Brown et al., 2008; Belyk and Brown, 2017; Eichert et al., 2020). While the ventral region (vLMC) is thought to support phonation and speech sound coordination (Eichert et al., 2020; Hickok et al., 2023), the dorsal region (dLMC) is more closely associated with pitch modulation (i.e., raising and lowering pitch; Dichter et al., 2018). An important question is therefore whether the LMC contributes to prosody perception and, if so, whether the dLMC contributes more strongly than the vLMC. Resolving this question would help determine whether effector-relevant motor area involvement represents an organizational principle that extends from segmental to suprasegmental features.

While previous studies have shown that PMC activity encodes categorical prosodic information (Tomasello et al., 2022; Baek et al., 2026) and that disrupting the PMC impairs prosody perception (Sammler et al., 2015), they have never tested whether these areas are involved in vocal pitch production, with one exception for lexical tones (Liang et al., 2023). Yet, because lexical tones and prosody serve distinct linguistic functions, it remains to be shown whether pitch motor areas contribute to prosodic category perception.

To address this gap, we acquired magnetoencephalography (MEG) data from participants during two sessions: First, they sang simple five-tone melodies to localize pitch motor areas, encompassing the putative vLMC and dLMC (Fig. 1A). Then, participants identified the prosody (Statement vs. Question) of single words varying in pitch contour on a five-level continuum to probe prosodic category perception (Fig. 1B). The prosody perception data were previously used for a different research question (Baek et al., 2026). If prosodic category perception recruits motor regions relevant to vocal pitch production, perceived prosodic categories should be decodable from the localized regions, with a potentially stronger contribution from the dLMC. Furthermore, if these neural patterns are behaviorally relevant to prosodic categorization, decoding performance should mirror perceptual discriminability, with lower decodability for ambiguous than clear prosody levels, before category judgments are typically reported, but not necessarily later. Finally, if pitch motor areas actively contribute to prosodic category processing, they should transfer category-related information to auditory regions rather than merely receive auditory input (e.g., Möttönen et al., 2013; Si et al., 2017).

**Figure 1.**
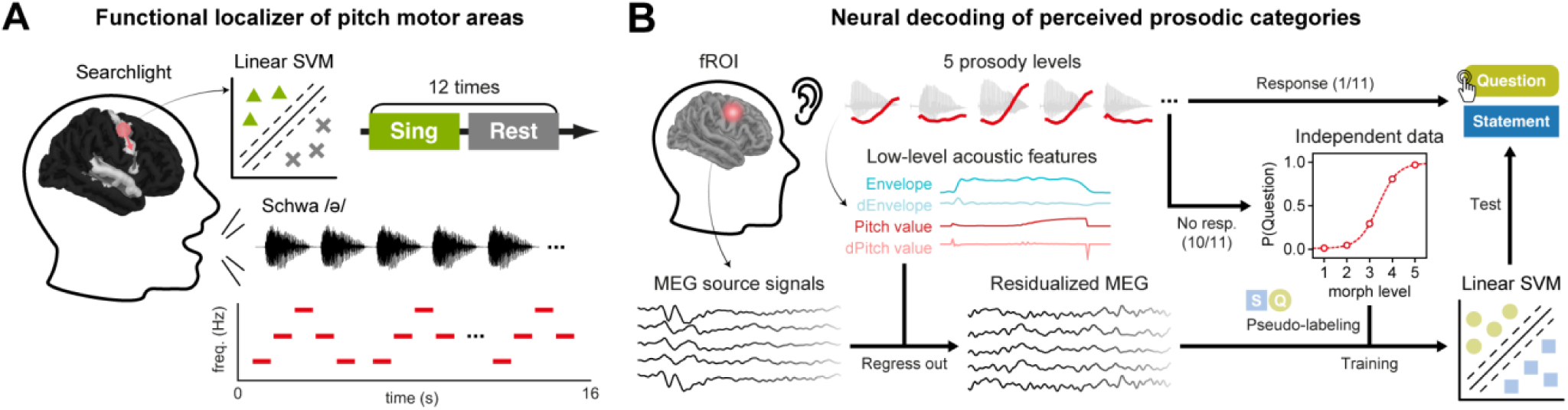
Experimental design and analysis workflow. **A**, Functional localizer of pitch motor areas. The localizer consisted of 12 singing and 12 resting trials (16 s each) in alternating order. Singing involved producing simple five-tone melodies with rising-falling contours on a neutral vowel (schwa /ə/) to engage the larynx through the production and modulation of vocal pitch while minimizing movement of supralaryngeal articulators. Source-reconstructed magnetoencephalography (MEG) data were analyzed using a searchlight-based (radius = 5 mm) linear support vector machine (SVM) to distinguish singing from resting trials. **B**, Neural decoding of perceived prosodic categories. In the main MEG experiment, participants identified the prosody (Statement vs. Question) of single words varying in pitch contour on a five-level continuum. Source MEG activity patterns within each functional region of interest (fROI) defined by **A** were first residualized for the modeled acoustic features of the stimuli (d, derivatives) to reduce contributions from low-level acoustic processing. The residualized MEG signals were then used to train and test linear SVMs in a time-resolved manner. Linear SVMs were trained on trials with pseudo-labels derived from independent behavioral data (10/11 of the trials used in the analysis). Decoding performance was tested on the remaining subset of trials (1/11) with prosody labels assigned by the participants.

## Materials and Methods

Part of the dataset collected in this study, including behavioral data and MEG recordings during the perception session, was previously analyzed by Baek et al. (2026) to address a separate research question: While Baek et al. (2026) tracked and compared the representational transfer of phonemes and prosody, from acoustics to categories along the cortical speech streams, and challenged serial hierarchical processing models of speech perception (e.g., Hickok and Poeppel, 2007; DeWitt and Rauschecker, 2012), the present study scrutinized the generalizability of motor theories of speech perception to prosody, i.e., to suprasegmental information produced by the larynx (source) rather than segmental information (phonemes) produced by the articulators (e.g., D’Ausilio et al., 2009; Möttönen and Watkins, 2009). To do so, we first localized motor regions relevant to vocal pitch production using an independent singing localizer, and then examined whether and how these regions relate to the behavioral categorization of words with varying prosodic pitch contours as statement or question. The stimuli, procedure, and data acquisition parameters of the perception session are described in detail in Baek et al. (2026).

### Participants

Thirty-four native German speakers were recruited for this study. All participants provided written informed consent prior to participation, and self-reported normal hearing and no history of neurological or psychiatric disorders. Following data collection, five participants were excluded from further analyses due to low behavioral performance (n = 4) or poor MEG data quality (n = 1). This resulted in a final sample of 29 participants (15 females), aged 26 ± 4 years (mean ± SD) and assessed as right-handed by the Edinburgh Handedness Inventory (median laterality quotient = 100, 10^th^–90^th^ percentiles = 80–100; Oldfield, 1971). Four participants (all males) did not take part in the singing session (see below). This study adhered to the ethical standards of the Declaration of Helsinki and was approved by the Ethics Committee of the Medical Faculty, Leipzig University, Germany (403/14-ff).

### Stimuli

Stimuli were single words that varied in 5 × 5 steps along two orthogonal dimensions: prosodic pitch contour and word-initial phoneme. The current study focused specifically on prosody, while the phoneme dimension was included for a separate study (Baek et al., 2026). The stimuli were constructed based on natural recordings of the German nouns “Bar” (pub) and “Paar” (pair), uttered as both statement and question by two native German speakers (one female). The recordings were made in a soundproof room using a Røde NT55 microphone and Audacity® (v2.0.5, sampling at 16 kHz, 16-bit, mono).

For each speaker, the recordings were morphed using MATLAB (R2014a, MathWorks) and STRAIGHT (Kawahara and Mutsui, 2003) to create 61 evenly spaced steps along two separate dimensions: pitch contour, ranging from statement (falling) to question (rising), and word-initial phoneme, ranging from /b/ to /p/. This procedure yielded 3,721 stimuli per speaker (7,442 stimuli in total), forming a 61 × 61 grid of pitch contour and word-initial phoneme. In particular, pitch contour morphing involved decomposing the word stems (i.e., excluding the word-initial phoneme) into five acoustic features (*F*_0_, frequency structure, duration, spectrotemporal density, and aperiodicity), followed by resynthesizing these features after interpolation in 2% steps from 0% (statement) to 120% of the original question.

From the 61 × 61 grid, 5 × 5 morph levels were selected for each speaker and tailored to individual participants based on a behavioral pretest using an adaptive staircase method (Kaernbach, 1991). The pretest aimed to identify five morph levels centered on individual category boundaries and to match the categorization difficulty across both dimensions and across speakers. Stimuli corresponding to these 5 × 5 morph levels were first selected in the behavioral experiment and then presented during the MEG perception session (see *Experimental design*). Stimulus duration ranged between 464 and 500 ms. The selected morph steps and their acoustic features are summarized in Table S1. Further detailed information about the phoneme morphing and the pretest procedure, as well as the selection of morph levels is provided in Baek et al. (2026).

### Experimental design

This study consisted of two parts: a behavioral and an MEG experiment. The behavioral experiment was designed to assess whether participants perceived prosody categorically and to validate the stimulus selection from the pretest (see *Stimuli*). To do so, participants were presented with their individualized set of stimuli, in three runs, each containing four blocks, i.e., two blocks for each speaker. For each speaker, participants were asked to identify the prosody (“Statement”/“Question”) in one block, or the word (“Bar”/“Paar”) in the other block. The two blocks of each speaker were presented back-to-back and contained the same stimuli in identical order. The task was indicated at the beginning of each block, and a fixation cross in the center of the screen remained visible throughout, with its color serving as a reminder for the current task (red for prosody, green for phonemes). In addition, the two corresponding response options were constantly displayed on the left and right sides of screen. The order of tasks and speakers within each run was randomized and counterbalanced across participants. Stimuli were delivered at a comfortable volume (60–70 dB SPL) via Beyerdynamic DT 770 PRO headphones in a soundproof chamber.

Every block comprised 56 trials, beginning with six practice trials utilizing clear morph levels (i.e., 1 or 5) of the respective task. The subsequent 50 trials featured 25 unique stimuli (5 × 5 morph levels) presented in a pseudo-randomized order, structured following the principles of a type-1 index-1 sequence to mitigate potential carry-over and position effects (Nonyane and Theobald, 2007). In each trial, participants identified the stimulus by pressing a button on a response box corresponding to the on-screen category labels. Button assignment was counterbalanced across participants. Responses were allowed up to 2500 ms after stimulus onset, immediately preceding the start of the next trial. The procedure was controlled by Presentation® software (v18.0, Neurobehavioral Systems) and lasted approximately one hour, with optional breaks provided between runs. For the purpose of the current study, data from the phoneme blocks were not included in the analyses.

The MEG experiment was conducted on a separate day, in an electromagnetically shielded room, comprising a singing session and a perception session. The singing session aimed to localize motor areas relevant to vocal pitch production, whereas the perception session investigated the involvement of these areas in the perception of prosodic categories. During the perception session, stimuli were delivered via air-conduction earphones (ER3-14A/B, Etymotic Research) at an average of 74 dB SPL. Intensity was adjusted for individual participants based on their stimulus-detection threshold, measured prior to the MEG experiment using a stimulus not presented later.

The singing session consisted of a single run, in which participants alternated between singing and resting trials (Fig. 1A). Before the session, participants were shown a visual representation of a five-tone rising-falling pitch contour (bottom left of Fig. 1A) and listened to a female experimenter (lay singer) singing the melody. They then practiced reproducing the overall contour comfortably within their natural vocal range. Singing on schwa /ə/ was chosen to engage the larynx through the production and modulation of vocal pitch while minimizing movements of supralaryngeal articulators. Additionally, participants were explicitly instructed to minimize not only movements of the articulators but also of face and head, such as widening their eyes or raising their head with increasing pitch.

During singing trials, a visual prompt “sing” indicated when participants should begin producing the practiced five-tone melody. They were asked to repeat the melody (in a loop) until a stop cue “rest” was presented on screen. During resting trials, participants were instructed to remain still and minimize their bodily movement. There were 12 singing and 12 resting trials, each lasting 16 s. To encourage natural vocal production, singing performance was not evaluated quantitatively. Yet, an experimenter monitored participants’ singing from outside the recording room and confirmed compliance with the task instructions.

The perception session consisted of six runs, each following a similar structure as the behavioral experiment in terms of stimulus randomization, block order, and counterbalancing of tasks and speakers. The only difference was the way behavioral responses were registered: In contrast to the behavioral experiment, a delayed response task with pseudorandom key assignment was used to isolate categorization-related activity from response-related neural signals. Moreover, overt behavioral responses were collected only in one-sixth of the trials to maximize the overall number of stimulus presentations within the session.

Participants were instructed to listen carefully and to categorize each stimulus in their mind while fixating on a central cross, whose color served as a task reminder (red for prosody, green for phonemes). In response trials, randomly interspersed within each block, response options showed up on the left and right sides of the screen. This prompted participants to indicate their response by pressing one of two buttons with their left or right thumb. Button assignment was randomized and balanced across stimuli, tasks, and speakers to prevent confounding with preparatory motor activity. The stimulus onset asynchrony was 2800 ms, jittered randomly between −500 and 500 ms. In response trials, an additional 2000-ms response window was provided, followed by a 1000-ms pause prior to the next stimulus. In the present study, analyses included all trials without responses (across both tasks) as well as all trials with prosody responses, resulting in a 1:11 response-to-stimulus ratio (Fig. 1B). Trials with phoneme responses were included only in a control analysis (see below).

The MEG experiment was controlled by Presentation® software (v18.0, Neurobehavioral Systems) and lasted up to 2.5 hours. Participants could take optional breaks between runs.

### Behavioral analysis

Behavioral data from the behavioral experiment and the MEG perception session were analyzed to assess the perception of prosodic categories. For each experiment and participant, we first calculated the proportion of “Question” responses across the five prosody levels for each speaker, collapsed across runs. Missed trials were excluded from the calculation. Four participants were excluded from further analyses because the difference between the lowest and highest proportions for the five morph levels (averaged across speakers) was less than 0.6 in at least one experiment (behavioral or MEG), suggesting that they had difficulties in perceiving clear “Statement” or “Question” categories.

The response proportions were then used to fit psychometric functions for each participant, experiment, and speaker following the formula below:

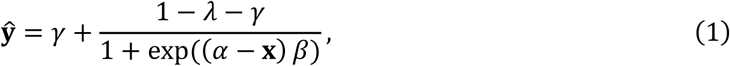

where **x** is the morph levels (normalized to 1–5), and **ŷ** is the estimated proportion. *α* indicates the point of subjective equality (PSE), where the function predicts equal proportions for both categories. *β* determines the slope at *α* and serves here as an index of perceptual difficulty. *γ* and *λ* represent the guess and lapse rates, corresponding to the lower and upper asymptotes of the function, respectively.

Curve fitting was performed using the `PAL_PFML_FIT` function (with @PAL_Logistic) from the Palamedes toolbox (v1.7.0) in MATLAB (R2014a, MathWorks) for the behavioral experiment, and the `curve_fit` function from the SciPy library (v1.16.3) in Python (v3.13.11) for the MEG perception session. In both implementations, *γ* and *λ* were fixed at 0.01. The fitted PSE and slope were separately compared between speakers and experiments at the group level using repeated measures analyses of variance (rmANOVAs). As no speaker-related effects were found for either measure (PSE: speaker, *F*_1,28_ = 0.29, *p* > 0.05, 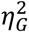 = 0.00; speaker × experiment, *F*_1,28_ = 3.89, *p* > 0.05, 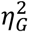 = 0.01; log slope: speaker, *F*_1,28_ = 0.80, *p* > 0.05, 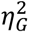 = 0.00; speaker × experiment, *F*_1,28_ = 0.32, *p* > 0.05, 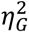 = 0.00), behavioral data were collapsed across speakers when reporting psychometric fits (Fig. 2) and conducting subsequent analyses, except for the pseudo-label generation described in *Neural decoding of perceived prosodic categories*.

**Figure 2.**
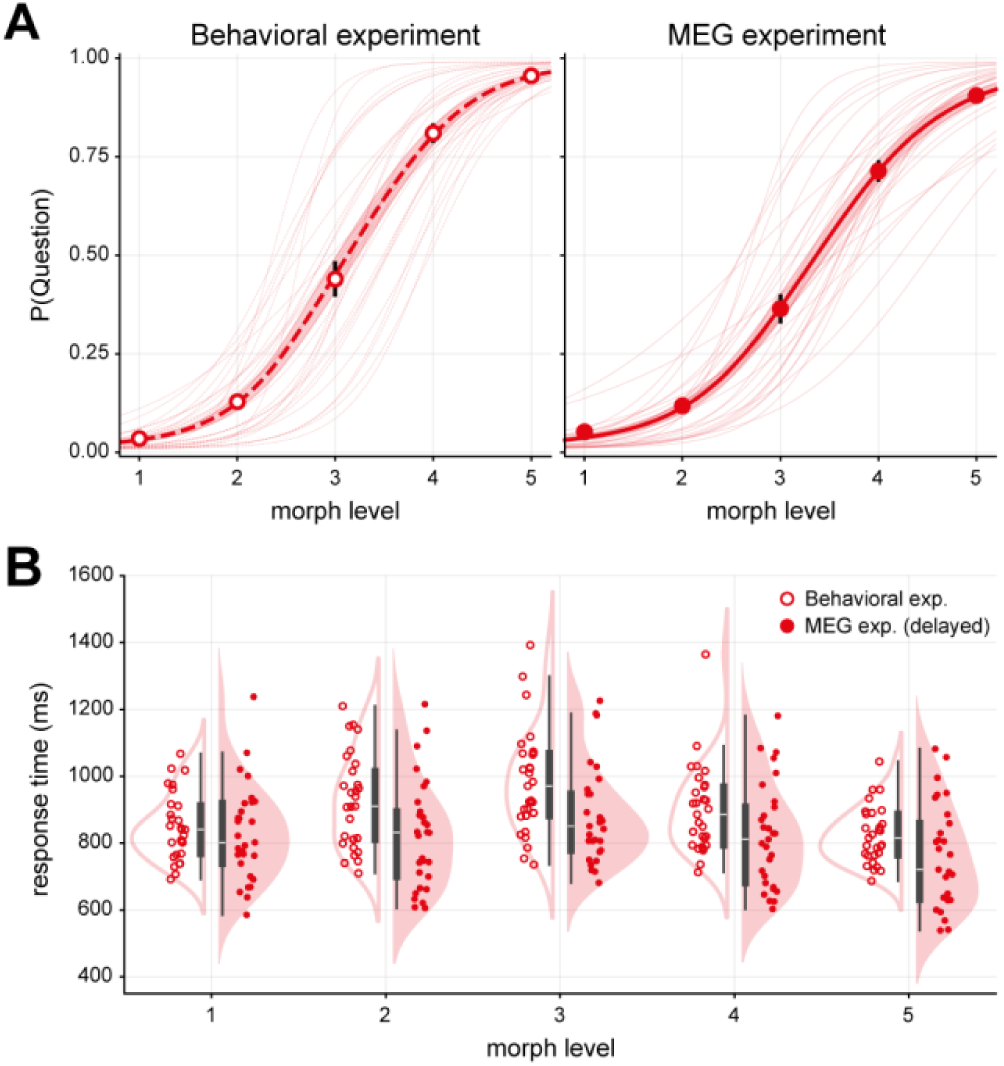
Behavioral results. **A**, Psychometric functions fitted to “Question” responses across five prosody morph levels in the behavioral (left) and the MEG (right) experiment. Shaded areas and error bars denote the standard error of the mean (SEM). Thin lines represent individual participant fits. **B**, Distributions of response times (RTs) across the five prosody levels for the behavioral (empty) and the MEG (filled) experiment. Boxplots represent the interquartile range (IQR) of RTs, with horizontal white lines indicating the median. Whiskers extend to 1.5*×* IQR. Individual participant medians are superimposed as jittered circles. Note that, in contrast to the behavioral experiment, the MEG experiment employed a delayed-response paradigm. RTs are time-locked to stimulus onset in the behavioral experiment and to the occurrence of the response prompt after stimulus offset in the MEG experiment.

Additionally, response times (RTs) were analyzed to further characterize the perception of prosodic categories. This analysis included all trials with an overt prosody response, as the stimuli varied gradually in pitch contour along the five-level continuum and therefore had no objectively correct or incorrect responses. For each experiment and participant, median RTs were computed across trials for each of the five morph levels, collapsed across speakers. Missed trials were again excluded.

### MEG data acquisition and preprocessing

During the MEG experiment, neuromagnetic activity was recorded using a 306-channel Neuromag Vectorview system (MEGIN), consisting of 102 magnetometers and 204 planar gradiometers. MEG data were sampled at 1 kHz, with online filtering from DC to 330 Hz. Throughout the recording, five head-position indicator coils were used to continuously monitor participants’ head positions. For participants who completed the singing session, additional 3–4 min empty-room recordings were collected before and after the MEG experiment using the same acquisition parameters.

The MEG data and empty-room recordings were preprocessed using MNE-Python (v1.11.0) in Python (v3.13.11). For each session and run, spatiotemporal signal-space separation (tSSS; internal/external component order = 11/2) was applied to suppress external interference, correct for head movements, and interpolate semi-automatically identified noisy channels (singing session: median = 5, 10^th^–90^th^ percentiles = 3–8; perception session: median = 6, 10^th^–90^th^ percentiles = 3–8). For empty-room recordings, the same parameters as in the corresponding singing session were used. Line noise (50 Hz) and its harmonics were then removed using the `dss_line` function (ZapLine; de Cheveigné, 2020) in the meegkit library (v0.1.9). Subsequently, signals were lowpass-filtered at 40 Hz (with a 10-Hz transition bandwidth) and highpass-filtered at 0.3 Hz (with a 0.3-Hz transition bandwidth) using separate one-pass, zero-phase, non-causal finite impulse response (FIR) filters (Hann window). In particular, the highpass-filter was designed to substitute the classical baseline correction (Widmann et al., 2015; Maess et al., 2016).

The filtered signals were downsampled to 200 Hz, followed by independent component analysis (ICA) to identify physiological artifacts, such as eye movements and heartbeats (Picard; Ablin et al., 2018). The ICA solution was computed separately for each participant and session using data concatenated across runs. To improve decomposition, the ICA solution was derived from data highpass-filtered at 1 Hz; these 1-Hz filtered signals were used solely for estimating the ICA weights and did not proceed to subsequent analyses. Based on the resulting ICA solution, artifactual components were identified semi-automatically and rejected (singing session: median = 6, 10^th^–90^th^ percentiles = 4–9; perception session: median = 9, 10^th^–90^th^ percentiles = 5–12). Empty-room recordings were cleaned by the same ICA solution and component rejection criteria as the corresponding singing session.

After the artifact rejection, the perception data were epoched from −200 to 1000 ms relative to stimulus onset, while the singing data were epoched from 0.5 to 15.5 s relative to trial onset to exclude potential carry-over from the preceding trial. Epoching was not applied to empty-room recordings. Epochs containing excessive noise were discarded by Autoreject (v0.4.3; Jas et al., 2017). Accordingly, one participant was excluded because more than 20% of trials were rejected in at least one session. For the remaining participants, a median of 24 epochs (10^th^–90^th^ percentiles = 23–24) for the singing session and 1,200 epochs (10^th^–90^th^ percentiles = 1,196–1,200) for the perception session were retained for further analyses.

### Source reconstruction

Source signals were reconstructed from the preprocessed MEG data using anatomical constraints derived from individual high-resolution T1-weighted images (1 × 1 × 1 mm³), acquired with a 3-T magnetic resonance imaging (MRI) scanner (Siemens). For each participant, anatomical images were processed with FreeSurfer (v7.3.2) to generate a volume conduction model and a cortical source space. The volume conduction model was estimated using a single-layer boundary element model based on the inner skull surface obtained via the watershed algorithm. The source space consisted of 10,242 vertices per hemisphere distributed across the cortical surface.

MEG and MRI data were co-registered by aligning fiducial points and digitized points on the head model using an iterative procedure. Sensor-level signals were then projected onto the source space using a linearly constrained minimum variance (LCMV) beamformer (Van Veen et al., 1997). Source orientations were optimized per location to maximize the output power and reduced to a scalar estimate under a unit-noise-gain constraint with a regularization parameter of 0.05 (e.g., Sekihara and Nagarajan, 2008; Westner et al., 2022). For the singing session, the LCMV spatial filter was computed using data covariance across all singing and resting epochs and noise covariance from empty-room recordings. For the perception session, data covariance was computed from the post-stimulus interval (0–1000 ms), and noise covariance from the prestimulus baseline (−200 to 0 ms). Covariance estimation was regularized by the Ledoit-Wolf method with cross-validation (Ledoit and Wolf, 2004), with rank adjusted to account for the degrees of freedom lost during tSSS and ICA preprocessing (singing session: median = 113, 10^th^–90^th^ percentiles = 108–117; perception session: median = 111, 10^th^–90^th^ percentiles = 107–116).

Source estimation was performed on a single-trial basis. For the singing session, the overall source power during individual singing and resting trials was estimated from their covariance (i.e., collapsing the time dimension), whereas for the perception session, time-resolved source activity was reconstructed for each trial. Finally, source estimates in individual native space were morphed into the common `fsaverage` space for subsequent analyses.

### Functional localization of pitch motor areas

Source data from the 25 participants who completed the singing session were used to localize pitch motor areas within the PMC (Fig. 1A). Specifically, a searchlight-based linear support vector machine (SVM) approach was employed to localize regions that discriminate between singing and resting, given its higher sensitivity relative to univariate methods (e.g., Haynes and Rees, 2006; Kriegeskorte et al., 2006). For each of the 25 participants, a linear SVM was trained and tested on multivariate source activity patterns within a cortical searchlight (5-mm radius) using nested leave-one-trial-out cross-validation. In the inner loop, the optimal margin parameter (*C*) was selected from a logarithmically spaced range (10^−5^ to 10^4^; 10 values) to maximize model sensitivity, and the resulting model was evaluated on the held-out trial in the outer loop. Input features were standardized based on the training data within each cross-validation fold, and classification accuracy was calculated by averaging across folds and assigned to the center vertex of each searchlight. This analysis was implemented using the `LinearSVC` function from the scikit-learn library (v1.7.2) in Python (v3.13.11).

Classification accuracy was then tested against chance (i.e., 0.5) at the group level across the 25 participants. Although accuracy maps were computed across the whole brain, statistical inference was restricted to vertices within three bilateral parcels defined by a combined version of a multimodal cortical parcellation (Glasser et al., 2016): the PMC, auditory cortex (AC), and superior temporal cortex (STC). The AC and STC were included for comparison as key components of the ventral auditory stream, which are also known to be sensitive to self-produced sounds (e.g., Hickok et al., 2011; Dichter et al., 2018).

Statistical significance was determined by one-sample, one-sided *t*-tests with a threshold of *p* < 0.001, Bonferroni-corrected jointly across all vertices within the six parcels. Although both hemispheres were included, we did not formulate or test a lateralization hypothesis because hemispheric differences were not a primary focus of the study.

As approximately 25% of vertices in both the left and right PMC were identified as significant (Table 1), a fixed number of 98 vertices—corresponding to 25% of the average vertex count across the bilateral PMC—was used to define equal-sized, group-level functional regions of interest (fROIs) in each parcel (the left and right PMC, AC, and STC). To promote spatial contiguity, a 5-mm FWHM Gaussian smoothing was first applied to the *t*-statistic map, after which the top 98 vertices in the smoothed *t*-statistics were selected within each parcel to define six fROIs (Fig. 3B). These group-level fROIs were then used in subsequent analyses of the MEG source data from the perception session (see below).

**Figure 3.**
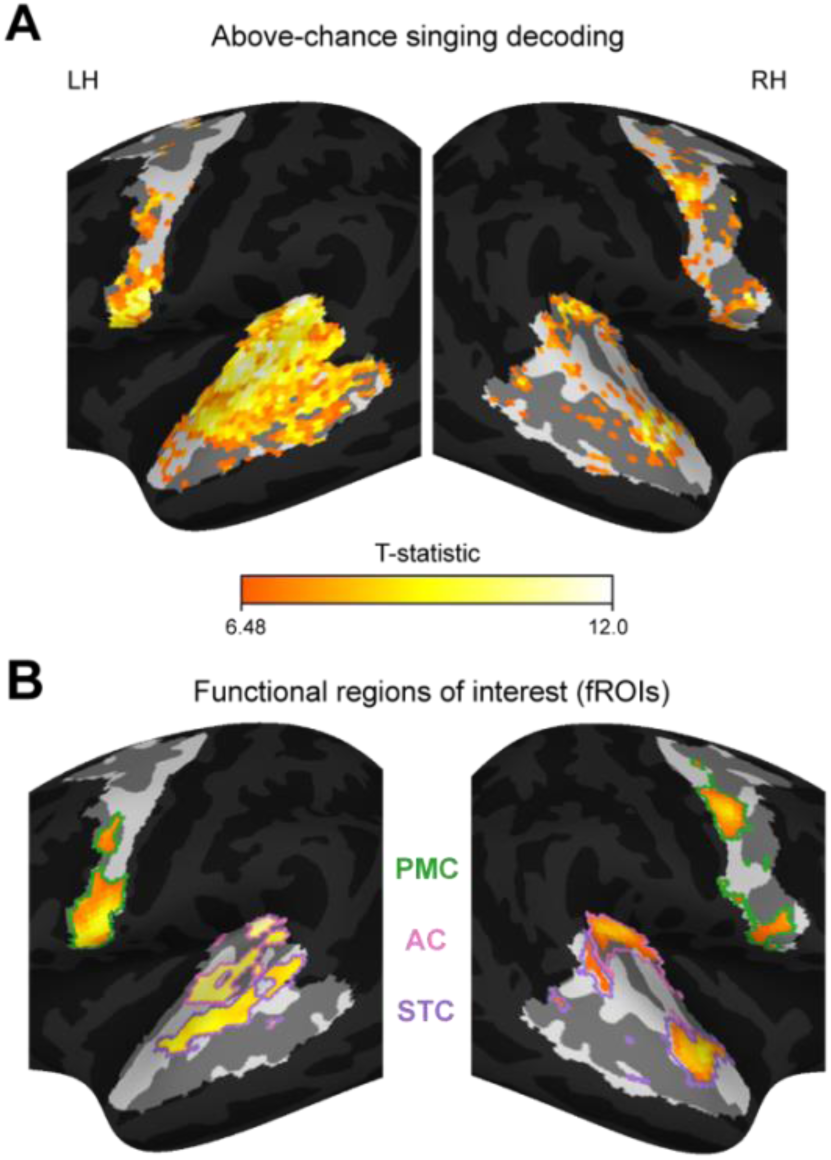
Above-chance singing decoding and fROIs. **A**, Group-level statistical map of vertices showing above-chance singing decoding (one-sample, one-sided *t*-tests against 0.5, *p* < 0.001 with Bonferroni correction across all vertices within unmasked regions) in the left (LH) and right (RH) hemispheres. **B**, fROIs defined by selecting a fixed number of 98 vertices ranked by Gaussian-smoothed (FWHM = 5 mm) *t*-statistics within the bilateral premotor cortex (PMC), auditory cortex (AC), and superior temporal cortex (STC).

**Table 1.** Statistics for above-chance singing decoding.

| **Region** | **Vertex count** | **%above-chance vertices** | **Average decoding performance (accuracy ± SEM, %)** | **Average *t*<sub>24</sub>-value** |
|---|---|---|---|---|
| L-PMC | 385 | 26.2 | 68 ± 2 | 5.64 |
| L-AC | 202 | 100 | 75 ± 2 | 9.33 |
| L-STC | 358 | 73.5 | 73 ± 2 | 7.35 |
| R-PMC | 398 | 25.9 | 67 ± 2 | 5.66 |
| R-AC | 165 | 34.6 | 67 ± 3 | 6.06 |
| R-STC | 370 | 25.7 | 68 ± 2 | 5.72 |

For each region, the total vertex count and the percentage of vertices showing above-chance decoding accuracy are reported. Average decoding performance and average *t*-values represent the grand mean across all significant vertices within each region. L-, left; R-, right; PMC, premotor cortex; AC, auditory cortex; STC, superior temporal cortex; SEM, standard error of the mean.

### Neural decoding of perceived prosodic categories

A second neural decoding analysis was performed within each of the six group-level fROIs on MEG source signals from the perception session for all 29 participants, to examine whether and when these areas contain information related to perceived prosodic categories (Fig. 1B), and how this compares between pitch motor (PMC) and auditory regions (AC and STC).

Motor theories assume that motor representations may provide category-related information that is not reducible to low-level acoustics (Liberman and Mattingly, 1985). We therefore tested whether perceived prosodic categories remain decodable after accounting for low-level acoustic responses. To this end, we used a temporal receptive field (TRF) model to predict neural responses to selected low-level acoustic features (see below) and regressed them out from the MEG source signals prior to the decoding analysis.

Four acoustic features were considered for TRF modelling: stimulus envelopes, pitch values (fundamental frequency, *F*_0_), and their respective derivatives. The envelopes were extracted as the magnitude of the Hilbert-transformed signal, and pitch values were estimated using Praat (v6.3.14; Boersma, 2001) at a 1-ms temporal resolution. Pitch values were further used to identify the pitch divergence point across the five prosody levels (see Fig. 4), serving here as a critical acoustic landmark for prosody. Details on pitch estimation and divergence point identification are provided in Baek et al. (2026). Stimulus envelopes and pitch values were lowpass-filtered at 40 Hz (using the same filter parameters as for the MEG data) and downsampled to 200 Hz. Their derivatives were then computed as first-order differences between successive samples. For samples in which pitch or its derivative was not defined, values were set to the 5^th^ percentile of the respective features for each speaker and participant. All acoustic features were padded with zeros to a uniform duration of 500 ms.

**Figure 4.**
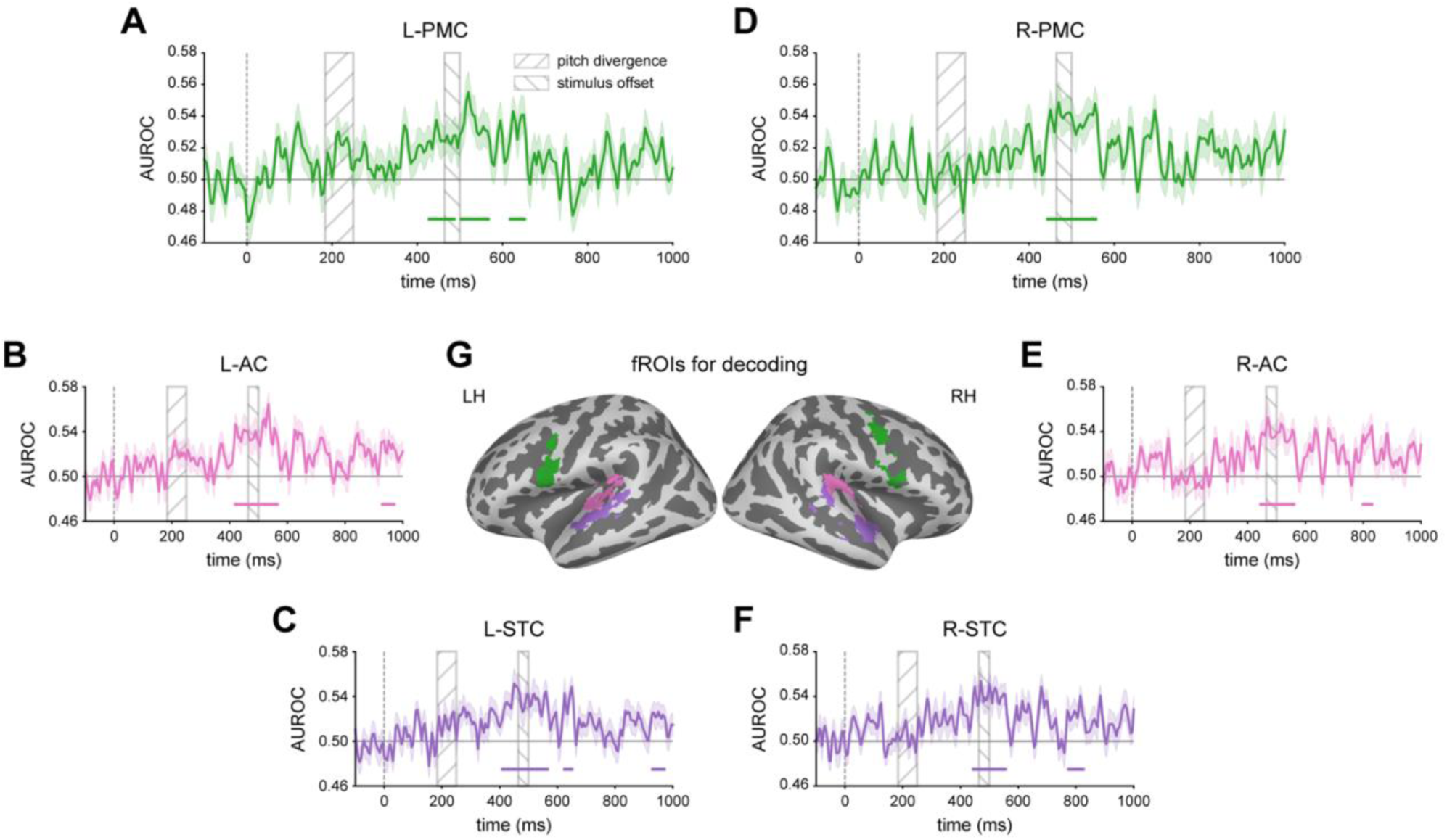
Neural decoding of perceived prosodic categories. Decoding models were trained on all trials without behavioral responses, using prosody pseudo-labels derived from independent behavioral data. The trained models were then tested on trials with overt responses from the prosody task. **A**-**C**, Time-resolved decoding performance, quantified as the area under the receiver operating characteristic curve (AUROC), in the left (L-) PMC (**A**), AC (**B**), and STC (**C**). Shaded areas indicate SEM. Horizontal lines above the x-axis denote significant time intervals (one-sample, one-sided *t*-tests against 0.5, cluster-forming threshold at *p*<0.05, family-wise error rate controlled at 0.05 across time points per fROI, fROI-wise Bonferroni-corrected *p*<0.0083). The rectangular areas with rising hatching (left) and falling hatching (right) represent the ranges of the pitch divergence point and stimulus offset, respectively. **D**-**F**, Corresponding decoding performance in the right (R-) PMC (**D**), AC (**E**), and STC (**F**). Statistical inference and figure details correspond to those provided in **A**-**C**. **G**, Locations of the six group-level fROIs used for decoding. Colors in the cortical maps correspond to those used for the decoding time series.

TRF models were fitted for each participant and fROI using the MNE-Python `ReceptiveField` function with 3-fold cross-validation. Response time lags ranged from 0 to 200 ms, and ridge regression was used as the estimator. For each fold, the optimal regularization hyperparameter was selected from a log-spaced range (10^−6^ to 10^6^; 13 values) based on the best predictive performance averaged across vertices. Prior to fitting, both acoustic features and MEG source signals were z-scored with respect to the training data. The predicted signals were then transformed back to the original scale and subtracted from the observed data to remove responses to low-level acoustics.

The residualized MEG signals were then used to decode perceived prosodic categories (Statement vs. Question) for each participant and fROI in a time-resolved manner. To this end, linear SVMs were trained at each time point using the `LinearSVC` function with a fixed margin parameter (*C* = 1). The models were trained on all trials without behavioral responses (995 ± 20 trials; mean ± SD), and tested on trials with prosody responses (96 ± 6 trials; mean ± SD).

For the training data, trial-wise prosody labels were generated from independent behavioral data. Specifically, labels were randomly assigned for each level and participant according to the estimated proportion of “Question” responses in the speaker-specific psychometric functions fitted in the behavioral experiment. The resulting pseudo-labels were balanced between the two prosodic categories across participants (Statement: 512 ± 49; Question: 484 ± 49 trials; mean ± SD; paired, two-sided *t*-test, *t*_28_ = 1.57, *p* > 0.05). After label assignment, the input neural data were normalized based on the training trials, and decoding performance on the test trials was quantified as the area under the receiver operating characteristic curve (AUROC).

Note that generating probabilistic pseudo-labels from independent data collected in a similar experimental paradigm is conceptually related to weak-supervision approaches commonly used in machine learning to handle missing labels. Although this procedure naturally induces noise into the training data, successful model evaluation with participant-labeled test data indicates that the classifier nevertheless captured meaningful neural patterns associated with perceived prosodic categories. To verify that decoding did not depend on a particular randomly generated set of prosody pseudo-labels, we regenerated the labels using 100 different random seeds and repeated the complete decoding analysis for each realization (Fig. S1).

As another control analysis, the prosody-trained classifiers were further evaluated on trials with phoneme responses (95 ± 6 trials; mean ± SD), to test whether the decoded neural patterns were specific to prosody (Fig. S2). Note that this test was not intended to assess motor contributions to phoneme perception. This would have required the localization of phoneme-related motor regions, such as lip motor areas for the labial phonemes /b/ and /p/, which was beyond the focus of the present study.

Lastly, the influence of task on the decoding of perceived prosodic categories was tested. To do so, linear SVMs were trained in the same way as described above, for each participant and fROI, but using only a subset of the training trials, either from the prosody or the phoneme identification task (see *Experimental design*). These task-specific models were then tested on the same held-out trials with prosody responses, and their performance was quantified as AUROC.

### Analysis of decoding importance scores

To characterize how different parts of the functionally localized pitch motor areas contributed to decoding perceived prosodic categories, we computed vertex-wise importance scores within each PMC fROI. The same procedure was also applied to the auditory fROIs. Because raw classifier weights are influenced by the covariance structure of the input features and cannot be interpreted directly as neural contributions, the weight vector for each participant and time point was transformed into an activation pattern following Haufe et al. (2014):

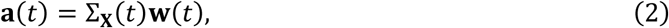

where Σ**_X_**(*t*) represents the covariance matrix of the training data from a given fROI, and **w**(*t*) is the weight vector of the corresponding linear SVM at time *t*. The resulting activation pattern, **a**(*t*), reflects how activity at each vertex covaries with the classifier decision variable. Vertex-wise importance scores were then derived as follows:

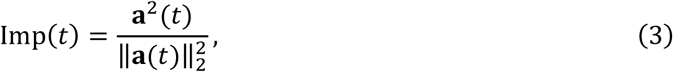

where squaring was performed element-wise and 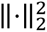 denotes the squared *L*_2_-norm. The resulting scores were nonnegative and summed to 1 across the 98 vertices within each fROI (see *Functional localization of pitch motor areas*), providing a variance-like measure of each vertex’s relative contribution to the multivariate pattern used to predict prosodic category labels. Accordingly, 1/98 ≈ 0.0102 represents uniform importance within an fROI and was used as a descriptive reference. For each participant and fROI, importance scores were averaged across significant decoding time points (Table 2).

**Table 2.** Cluster statistics for significant decoding of perceived prosodic categories.

| **fROI** | **Duration (ms)** | **Bonferroni<br>*p*-value** | **Average decoding performance<br>(AUROC ± SEM)** | **Average<br>*t*<sub>28</sub>-value** |
|---|---|---|---|---|
| L-PMC | 425-490 | 0.007 | 0.526 ± 0.006 | 2.40 |
|  | 500-570 | 0.002 | 0.536 ± 0.006 | 2.96 |
|  | 615-655 | 0.023 | 0.533 ± 0.008 | 2.88 |
| L-AC | 415-570 | <0.001 | 0.538 ± 0.005 | 3.45 |
|  | 925-975 | 0.017 | 0.527 ± 0.007 | 2.64 |
| L-STC | 405-570 | <0.001 | 0.535 ± 0.005 | 3.09 |
|  | 620-655 | 0.036 | 0.536 ± 0.008 | 3.17 |
|  | 925-975 | 0.040 | 0.522 ± 0.006 | 2.26 |
| R-PMC | 440-560 | <0.001 | 0.538 ± 0.005 | 3.46 |
| R-AC | 440-565 | <0.001 | 0.538 ± 0.006 | 3.29 |
|  | 795-835 | 0.036 | 0.535 ± 0.008 | 3.03 |
| R-STC | 440-560 | <0.001 | 0.540 ± 0.007 | 3.39 |
|  | 770-830 | 0.032 | 0.525 ± 0.007 | 2.29 |

For each functional region of interest (fROI), statistics for significant temporal clusters are reported. Average decoding performance and average *t*-values represent the grand mean across time points within each significant cluster. L-, left; R-, right; PMC, premotor cortex; AC, auditory cortex; STC, superior temporal cortex; AUROC, area under the receiver operating characteristic curve; SEM, standard error of the mean.

To compare importance scores between the ventral and dorsal PMC subregions, cortical surface patches were defined independently of the importance scores, based on the group-level smoothed *t*-statistic map from the singing localizer (Fig. 3B). Within each PMC fROI, we identified the two major spatial clusters corresponding to the ventral and dorsal subregions and selected the vertex with the highest smoothed *t*-value in each cluster. Each cortical patch comprised this peak and its neighboring vertices within a 5-mm radius, forming approximately size-matched ventral and dorsal subregions (8–13 vertices; Fig. 5A). For each participant and hemisphere, importance scores were averaged across vertices within each patch. Ventral-dorsal differences were then assessed separately for the left and right PMC using paired, two-sided *t*-tests.

**Figure 5.**
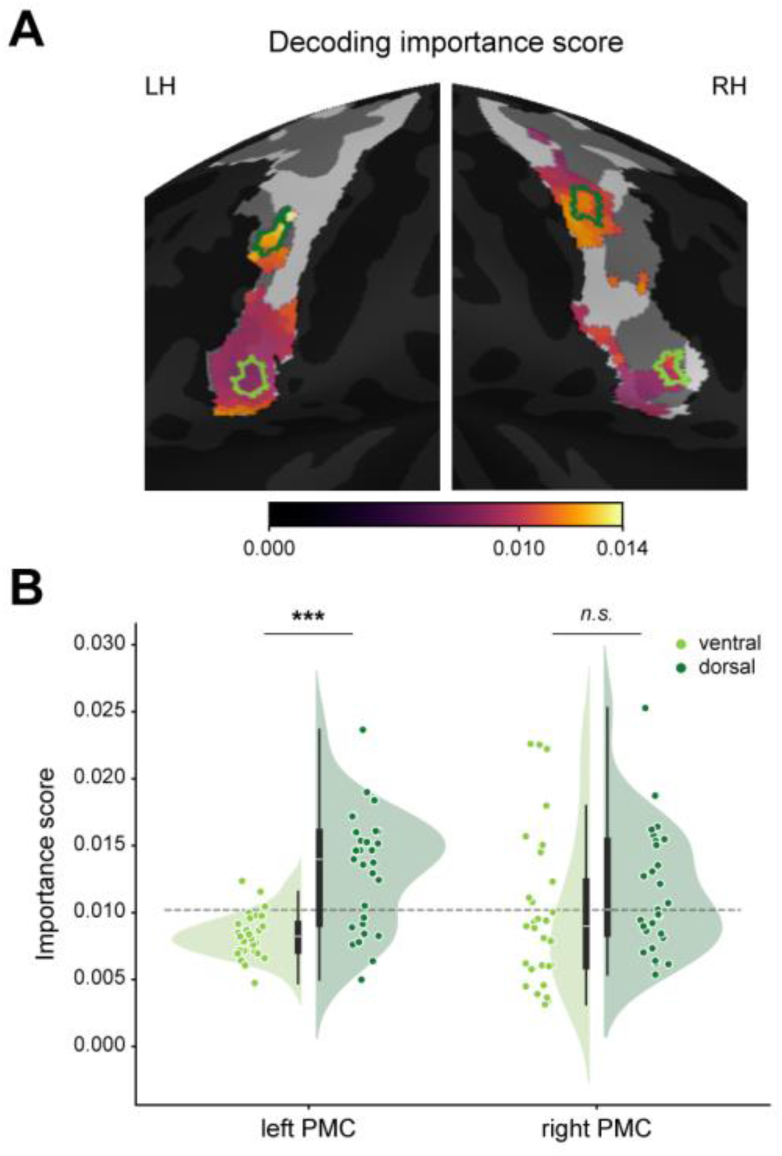
Decoding importance scores within the PMC. **A**, Group-averaged vertex-wise importance scores within the left and right PMC fROIs. Light and dark green outlines indicate the ventral and dorsal surface patches, respectively, used for statistical inference in **B**. Each patch comprised vertices within a 5-mm radius of the corresponding peak in the independent singing localizer map (Fig. 3B). **B**, Importance score distributions. Boxplots represent the IQR, with horizontal white lines indicating the median and whiskers extending to 1.5 × IQR. Dots denote individual participants. The dashed gray line indicates uniform importance of all vertices within each fROI (i.e., 1/98 vertices). Importance scores were significantly higher in the dorsal than in the ventral subregions of the left PMC, but not of the right PMC (paired, two-sided *t*-tests). *p* < 0.001, ***; not significant, *n.s*.

### Correlation between neural decoding and perceptual discriminability

To further investigate whether the neural decoding performance was related to the perceptual discriminability of the five prosody levels or to other processes not directly involved in prosodic categorization, such as the maintenance of category-related information, the linear SVMs previously trained for each participant and fROI across *all* levels (see above), were now evaluated separately for each level in the test set. Here, decoding performance was measured as accuracy because AUROC was not always defined—for instance, in cases where participants reported only a single category for a given prosody level. Level-wise decoding accuracy was then averaged across time points within each significant decoding interval (Table 2) for each participant and fROI. Intervals separated by only one sample (i.e., 5 ms) within a given fROI were aggregated prior to averaging.

Perceptual discriminability (*d*^′^) was derived from the psychometric function fitted to behavioral responses from the MEG experiment (Fig. 6A). In signal detection theory, *d*^′^ is defined as follows:

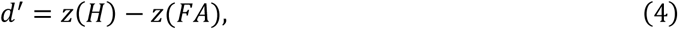

where *z*(·) denotes the inverse cumulative Gaussian distribution, and *H* and *FA* represent hit and false alarm rates, respectively. Hit rates were calculated directly from the behavioral responses (collapsed across speakers), whereas false alarm rates were estimated by re-fitting a psychometric function for each participant, including the estimation of guess (*γ*) and lapse (*λ*) rates (Eq. 1). The fitted guess and lapse rates served as baseline false alarm rates for “Question” and “Statement” responses, respectively (e.g., Wichmann and Hill, 2001; Kingdom and Prins, 2016). In addition, the estimated PSE (*μ*) was used to determine the target prosodic category for each level. Specifically, levels below the PSE were assigned “Statement”, while those above were assigned “Question”. Level-wise perceptual discriminability was then computed using Eq. 4, with log-linear correction (Hautus, 1995).

**Figure 6.**
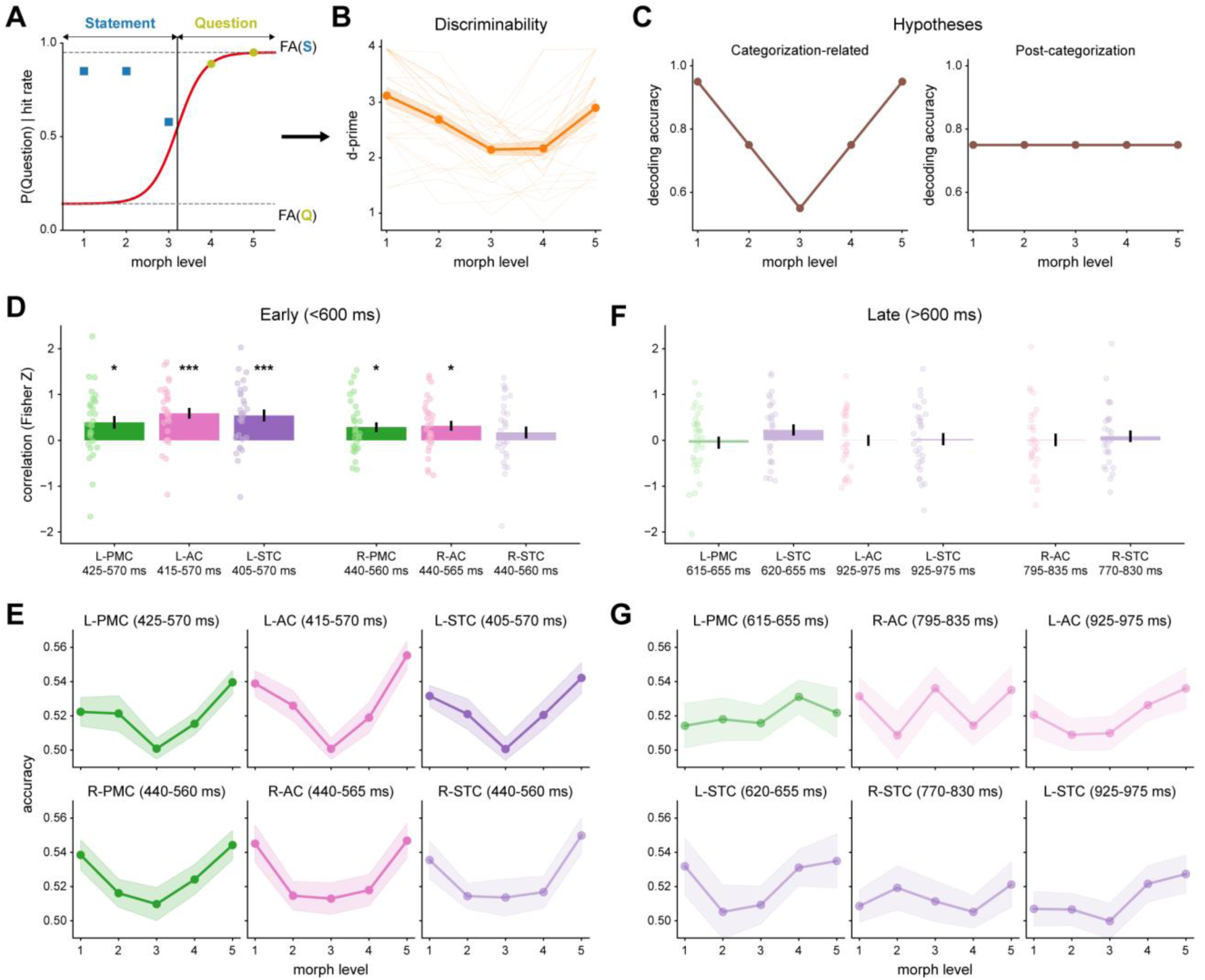
Correlation analysis between decoding accuracy and perceptual discriminability across prosody levels. **A**, Schematic of perceptual discriminability estimation (*d*′). Discriminability was derived from psychometric functions fitted to behavioral responses from the MEG experiment. The estimated upper and lower asymptotes were used as baseline false alarm rates (FA) for “Statement” (blue) and “Question” (yellow), respectively. Hit rates were calculated directly from behavioral responses, with the target prosodic category for each level determined by the estimated point of subjective equality (i.e., 50%-point). **B**, Estimated perceptual discriminability across the five prosody levels. Shaded areas indicate SEM. Thin lines represent individual participants. **C**, Hypotheses. If neural decoding reflects functional relevance to prosodic categorization, the decoding profile should mirror perceptual discriminability (left). By contrast, if decoding is related to later processes after category formation, the profile should be relatively constant across levels (right). **D**,**F**, Correlation (Fisher-Z transformed) between level-wise decoding accuracy and perceptual discriminability across fROIs within significant decoding intervals (see Fig. 4) before (**D**) and after (**F**) 600 ms relative to stimulus onset. Significant correlations are denoted by vivid colors; otherwise, pale colors (one-sample, one-sided *t*-tests against zero, false discovery rate (FDR) corrected *q*<0.05, *; *q*<0.001, ***). Error bars indicate SEM. Circles represent individual participants. **E**,**G**, Level-wise decoding accuracy within the significant intervals before (**E**) and after (**G**) 600 ms. Shaded areas indicate SEM.

Finally, decoding accuracy was correlated with perceptual discriminability across the five prosody levels for each participant, separately for each fROI and each significant time interval. This analysis tested the hypothesis that, if a region is associated with prosodic categorization at a given time, its level-wise decoding accuracy should mirror perceptual discriminability, resulting in a high correlation. By contrast, decoding related to later processes after category formation should remain relatively constant across levels, yielding a lower correlation (Fig. 6C). The correlations were quantified as Pearson’s correlation coefficient, followed by Fisher-Z transformation for group-level statistical testing.

### Multivariate transfer entropy (mTE) analysis

Lastly, directed information transfer between the fROIs was examined to test whether the pitch motor areas influence auditory regions during prosodic category processing. To do so, activation patterns were derived for each participant and fROI from the linear SVMs (via Eq. 2) starting at 400 ms post-onset, when perceived prosodic categories were significantly decoded (Table 2). The dimensionality of these activation patterns was then reduced using principal component analysis (PCA) for noise suppression and computational efficiency, retaining the components that explained 95% of the total variance (2–6 PCs). Importantly, the first principal component (PC1; explaining 68 ± 11%; mean ± SD) was adaptively excluded from the analysis if its cross-correlation between a given fROI pair peaked at zero-lag, to mitigate spurious effects driven by high spatial correlation or instantaneous shared variance.

Based on these dimensionality-reduced signals, directed information transfer between fROI pairs was measured using a recently developed mTE framework (Yu et al., 2019; Zhou et al., 2022). This framework combines Rényi’s *α*-order entropy—a generalization of Shannon entropy—with a Gaussian kernel trick, enabling the assessment of transfer entropy between two multivariate time series without estimating their individual or joint probability density functions. Instead, this approach relies on the normalized Gram matrix of the multivariate time series and uses its spectral information (i.e., eigenvalues) to estimate transfer entropy. The mathematical formalization and theoretical justification for this framework are detailed in Yu et al. (2020) and Zhou et al. (2022).

For mTE estimation, the current study followed the protocol described in Baek et al. (2026). However, while Baek et al. (2026) utilized nominal time lags, time-lagged signals were modeled here by phase space reconstruction (PSR; e.g., Wibral et al., 2011; Zhou et al., 2022). This technique expands the observed time series into a higher-dimensional space using time-delayed embedding, thereby capturing the latent dynamics of the underlying neural system (Takens, 1981). PSR was tailored to each fROI pair by defining a unified time delay to align their temporal dynamics. This delay was set as the minimum of the two fROI-specific delays, each estimated as the mode across its PCs based on average mutual information (Wallot and Mønster, 2018). Given this unified delay, embedding dimensions for the two fROIs were separately determined by the false nearest neighbor method (Wallot and Mønster, 2018). These parameters were then used to reconstruct time-delayed embeddings of category-related activation patterns, after which the directed connection from the source to the target fROI was quantified.

mTE was estimated for each participant and each fROI pair, with individual fROIs alternatively serving as source and target. However, these estimates were non-negatively biased and varied in scale across participants and directed connections (Boba et al., 2015), complicating comparisons and group-level statistical inference. To address this, mTE was normalized relative to a null distribution. The null distribution was generated by cyclically time-shifting the activation patterns of the source fROI across all possible offsets within the 5^th^ and 95^th^ percentile of the analyzed time window (i.e., 400–1000 ms post-onset), followed by embedding using the same PSR parameters as the original data. The mean and standard deviation of this distribution were then used to z-score the observed mTE, yielding a normalized value for each participant and directed connection.

### Statistical analysis

Group-level inference of perceived prosodic category decoding was performed using cluster-based permutation tests (Maris and Oostenveld, 2007). Temporal clusters were identified with one-sample, one-sided *t*-tests against chance (0.5) at *p* < 0.05, and the family-wise error rate was controlled at *p* < 0.05 across time points within each fROI. To compare decoding performance between the two task-specific models, one-sided instead of two-sided *t*-tests were used, reflecting the hypothesis that decoding would be higher during the prosody than the phoneme task. Permutation distributions were constructed by randomly shuffling decoding values relative to chance or task labels 10,000 times. Multiple comparisons across fROIs were controlled using a Bonferroni-corrected threshold of *p* < 0.0083 (i.e., 0.05/6 fROIs). For inference on decoding performance for each task-specific model, the threshold was adjusted to a Bonferroni-corrected threshold of *p* < 0.0042 to account for testing the same dataset twice.

The correlation between decoding accuracy and perceptual discriminability was evaluated at the group-level using one-sample, one-sided *t*-tests, based on the hypothesis that pitch motor areas are involved in perceptual categorization (Liang et al., 2023). Similarly, mTE of category-related activation patterns was tested against zero using one-sample, one-sided *t*-tests, motivated by prior evidence of auditory-motor interactions (e.g., Möttönen et al., 2013; Si et al., 2017). Multiple comparisons across correlations or directed connections were controlled using a false discovery rate (FDR) of *q* < 0.05.

### Code accessibility

The scripts used for the analyses and figure generation, as well as the processed behavioral and MEG data are publicly available in the following repository: https://osf.io/4u5kx/

## Results

### Perception of prosodic categories

Behavioral responses from both the behavioral and MEG experiments followed a typical psychometric function across the five prosody levels (Fig. 2A), indicating that participants perceived the stimuli categorically in both experiments. The response profiles in the two experiments were similar, with differences observed: The fitted PSE was higher in the MEG (3.4 ± 0.5) than in the behavioral experiment (3.1 ± 0.5; mean ± SD; paired, two-sided *t*-test, *t*_28_ = 3.98, *p* < 0.001), reflecting a stronger bias toward “Statement” over “Question” responses in the MEG experiment. Moreover, the fitted slope was shallower in the MEG experiment (log slope; paired, two-sided *t*-test, *t*_28_ = −2.82, *p* < 0.001), suggesting that participants found it more difficult to identify prosodic categories in the MEG experiment. These differences were likely driven by the different settings of the MEG experiment, such as the delayed response and variable response button assignments.

RTs further supported the prosodic category perception, featuring a characteristic inverted U-shape across the five prosody levels in both the behavioral and MEG experiments (Fig. 2B). This pattern indicates faster categorization for clear, prototypical prosody levels than for ambiguous ones, as supported by a significant main effect of level (*F*_4,112_ = 26.65, *p* < 0.001, 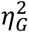 = 0.10), revealed by an rmANOVA comparing RTs across experiments and levels. RTs also differed significantly between the two experiments (*F*_1,28_ = 5.79, *p* = 0.023, 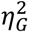 = 0.07), along with a significant interaction between experiment and level (*F*_4,112_ = 4.29, *p* = 0.006, 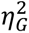 = 0.01). While RTs were measured from stimulus onset in the behavioral experiment, they were measured from the onset of the response options presented after stimulus offset in the MEG experiment. This delayed response requirement likely contributed to faster RTs and a less pronounced inverted U-shape in the MEG experiment. In contrast, responses in the behavioral experiment were generally made after 600-ms relative to stimulus onset, implying that prosodic categorization occurred prior to this time point.

Overall, these results demonstrate that speech prosody was categorically perceived, consistently across both the behavioral and the MEG experiment, with slight differences in response patterns attributable to experiment-specific constraints.

### Pitch motor areas in the premotor cortex

To functionally localize motor regions relevant to vocal pitch production, we applied a searchlight-based linear SVM approach to the MEG singing data to identify PMC subregions that distinguish singing from resting (Fig. 1A). This approach identified approximately 25% of the vertices within both the left and right PMC as showing above-chance singing decoding (Table 1). These vertices were largely clustered in the ventral and middle regions of the PMC, bilaterally (Fig. 3A). The ventral cluster was localized posterior to the inferior frontal gyrus, whereas the middle cluster was situated around the precentral gyrus, posterior to the middle frontal gyrus. These regions anatomically correspond to the vLMC and dLMC reported in previous studies (e.g., Brown et al., 2008; Dichter et al., 2018; Lu et al., 2023), supporting their relevance to vocal pitch production. However, this contrast may also capture broader processes associated with singing, such as respiration and vocal effort (e.g., Zarate, 2013; Herbst, 2017).

Singing was also decodable in the bilateral AC and STC, including Heschl’s gyrus and the planum temporale, as well as the superior temporal gyrus and sulcus (Fig. 3A). These findings likely reflect the neural responses of auditory regions to self-produced sounds (e.g., Hickok et al., 2011; Dichter et al., 2018). Interestingly, above-chance singing decoding was spatially more extensive and robust in the left AC and STC compared to their right-hemispheric homologues (see also Table 1), which may relate to left-hemispheric predominance in sensorimotor integration for vocalization (Hickok and Poeppel, 2007; Hickok et al., 2011; Behroozmand et al., 2015).

Building on these results, we delineated six fROIs, within the bilateral PMC, as well as the bilateral AC and STC for comparison. This involved first smoothing the statistical map to improve spatial contiguity and then deriving equally sized, group-level fROIs within each region (see Materials and Methods). As shown in Figure 3B, the localized pitch motor areas within the bilateral PMC comprised primarily two subregions, corresponding to the putative vLMC and dLMC. Notably, the dorsal area was more spatially pronounced in the right PMC, whereas the ventral area was more extensive in the left PMC. These asymmetries might reflect more dominant roles of the right dLMC in pitch control and the left vLMC in phonation (e.g., Hickok et al., 2011, 2023; Dichter et al., 2018).

Altogether, these findings suggest that the functional localizer identified PMC regions relevant to vocal pitch production, encompassing the putative vLMC and dLMC.

### Decoding perceived prosodic categories in pitch motor areas

Having identified pitch motor areas, we performed a time-resolved neural decoding analysis on the MEG perception data to determine whether and when these regions differentiate between the two prosodic categories. MEG source signals were first residualized with respect to the modeled acoustic features of the stimuli to reduce contributions from low-level acoustic processing and were then used to train classifiers with pseudo-labels generated from independent behavioral data (Fig. 1B; see Materials and Methods). The decoding time courses were consistent across different realizations of the prosody pseudo-labels (Fig. S1), indicating that the results were robust to their probabilistic generation. Furthermore, in another control analysis, the trained classifiers did not significantly discriminate phoneme categories (Fig. S2; all cluster *ps* > 0.05), demonstrating that the decoded neural patterns were specific to prosody.

Figure 4 shows the time-resolved decoding of perceived prosodic categories across the fROIs (see also Table 2). In the bilateral PMC fROIs (Fig. 4A,D), prosodic categories were decodable as early as ∼430 ms after stimulus onset, corresponding to approximately 200 ms after the pitch divergence point (184–250 ms), at which pitch contours began to clearly diverge across the five prosody levels. Decodability in these regions persisted beyond stimulus offset (464–500 ms), with an additional later time window in the left PMC fROI around 630 ms post-onset. These effects are unlikely to be explained solely by neural responses to the modeled low-level acoustic features, which were regressed out from the MEG source signals. Indeed, without MEG residualization, decoding performance was reduced and significant for a shorter duration before and around stimulus offset (Fig. S3), suggesting that responses captured by the acoustic model may instead partially obscure neural patterns associated with perceived prosodic categories.

Next, we computed vertex-wise importance scores to characterize the relative contributions of dorsal and ventral parts of the localized pitch motor areas to prosodic category decoding. In both the left and right PMC fROIs, the dorsal subregion generally showed greater decoding importance than the ventral subregion (Fig. 5A), with the highest scores located in the putative dLMC (left PMC: 0.014 ± 0.005; right PMC: 0.013 ± 0.005; mean ± SD). To test this apparent dorsal predominance, we defined ventral and dorsal cortical patches independently of the importance scores, based on the singing localizer (see Materials and Methods) and compared their importance scores (paired, two-sided *t*-tests). Importance scores were significantly higher in the dorsal than in the ventral patch of the left PMC (*t*_28_ = 4.83, *p* < 0.001), whereas the difference was not significant in the right PMC (*t*_28_ = 0.95, *p* = 0.35; Fig. 5B). The relatively stronger dorsal contribution, particularly in the left PMC, links prosodic category decoding to a motor subregion more closely associated with pitch modulation (e.g., Dichter et al., 2018; Finkel et al., 2019).

Perceived prosodic categories were also decodable in auditory regions (Fig. 4B,C,E,F; Table 2), in line with recent evidence of categorical prosodic representations in the AC and STC (Gnanateja et al., 2025; Baek et al., 2026). Similar to the PMC, significant decoding was observed starting at ∼430 ms post-onset and lasted beyond stimulus offset, consistent with parallel categorical prosody processing across spatially distributed cortical regions (Keshishian et al., 2023; Baek et al., 2026). Notably, the bilateral AC and STC also exhibited significant decoding in late time windows (within 800–1000 ms), which may reflect the maintenance or reinstantiation of category-related prosodic information due to the delayed response task. Within these regions, decoding importance was concentrated largely in lateral areas of the AC and posterior subregions of the STC (Fig. S4).

Lastly, we conducted follow-up analyses separately for the prosody and phoneme tasks to investigate how task demands influenced the decoding of perceived prosodic categories (see Materials and Methods). Although direct comparisons between the two task-specific models did not yield significant modulation effects (all cluster *ps* > 0.05), separate evaluations of each model revealed differences in the decoding patterns. Specifically, significant decoding within 400–600 ms post-onset was replicated in all fROIs in the prosody task, whereas in the phoneme task, the effect was found only in the right PMC fROI and for a shorter duration (Fig. S5; Table S2). These findings indicate that category-related prosodic representations are generally more robust during the relevant task, alongside task-invariant representations in the right PMC.

Taken together, these results highlight that, beyond auditory regions, motor areas relevant to vocal pitch production contained information about perceived prosodic categories, suggesting effector-relevant motor area involvement. Within these areas, the dorsal subregion, which is associated more closely with pitch modulation, contributed more strongly to decoding, particularly in the left PMC. Moreover, the task-invariant category-related prosodic information in the right PMC point to its relatively automatic involvement, which may provide a fundamental scaffold for prosodic category processing (cf. Chevillet et al., 2013, for phonemes).

### Relevance of pitch motor areas to perceptual categorization

While neural decoding has demonstrated that pitch motor areas contain information about perceived prosodic categories, the functional relevance of this information to perceptual categorization remains to be established. We therefore performed a correlation analysis to relate neural decoding performance to perceptual discriminability (*d*^′^) across the five prosody levels. Perceptual discriminability was derived from psychometric functions fitted to behavioral responses from the MEG experiment, including estimates of guess and lapse rates (Fig. 6A; see Materials and Methods). As shown in Figure 6B, perceptual discriminability between two prosodic categories exhibited a typical U-shaped curve, reflecting the increased difficulty in categorizing ambiguous prosody levels compared to clear, prototypical ones.

The estimated discriminability was then correlated with level-wise decoding accuracy, obtained by re-evaluating the classifiers (trained across all levels) separately for each prosody level. This re-evaluation was carried out within the fROIs and time intervals that previously showed significant decoding (Fig. 4; Table 2). This analysis tested the hypothesis that, only in fROIs and time points that are related to perceptual categorization, level-wise decoding accuracy should show a similar U-shaped profile and therefore result in high correlations; otherwise, the decoding profile should remain relatively flat across levels (Fig. 6C).

The correlation analysis revealed that decoding accuracy in both the left and right PMC fROIs significantly mirrored perceptual discriminability. Crucially, this correlation was confined to the time intervals earlier than 600 ms relative to stimulus onset (Fig. 6D; left PMC: *r* = 0.29 ± 0.09, *t*_28_ = 2.80, FDR-corrected *q* = 0.014; right PMC: *r* = 0.21 ± 0.08, *t*_28_ = 2.65, FDR-corrected *q* = 0.016; mean ± standard error [SEM]). These intervals encompassed stimulus offset and preceded the period in which participants generally made behavioral responses (Fig. 2B), suggesting that during these time points, pitch motor areas were functionally related relevant to perceptual categorization. Indeed, level-wise decoding accuracy in the bilateral PMC fROIs exhibited a characteristic U-shaped profile before 600 ms (Fig. 6E), indicating that decoding was more challenging for perceptually ambiguous stimuli than for more prototypical prosody levels. Notably, no such relationship was found after 600 ms (Fig. 6F; left PMC: *r* = −0.01 ± 0.10, *t*_28_ = −0.37, FDR-corrected *q* > 0.05; mean ± SEM), where the decoding profile became relatively constant (Fig. 6G). This suggests that decoding during this later time window may instead reflect other processes after categorization, such as the maintenance of category-related prosodic representations driven by the delayed response requirement.

Furthermore, decoding accuracy in auditory regions, including the bilateral AC and left STC, followed the pattern of perceptual discriminability (Fig. 6D,E), suggesting their co-involvement in perceptual categorization. Similarly, significant correlations were observed only during the intervals prior to 600 ms post-onset (left AC: *r* = 0.44 ± 0.08, *t*_28_ = 4.97, FDR-corrected *q* < 0.001; left STC: *r* = 0.39 ± 0.09, *t*_28_ = 4.16, FDR-corrected *q* < 0.001; right AC: *r* = 0.24 ± 0.08, *t*_28_ = 2.95, FDR-corrected *q* = 0.013; mean ± SEM), which may represent a critical time window for the categorization process, at least within the context of our experimental design. In contrast, no significant correlations were observed in later time windows (Fig. 6F,G; FDR-corrected *qs* > 0.05), potentially reflecting the maintenance or reinstantiation of category-related prosodic representations.

Again, the observed correlations are not likely to be driven simply by neural responses to the modeled low-level acoustic features, as responses predicted by these features were removed from the MEG source signals. Moreover, these correlations were no longer significant when the same analysis was performed without MEG residualization (all FDR-corrected *qs* > 0.05; Fig. S6), suggesting that responses captured by the acoustic model may instead obscure the relationship between neural and perceptual discriminability.

Overall, these findings underscore that pitch motor areas, alongside auditory regions, were functionally relevant to perceptual categorization during specific time windows prior to behavioral responses, rather than merely carrying category-related prosodic information.

### Category-related information transfer between pitch motor and auditory areas

Despite significant decoding observed in the PMC fROIs and its functional relevance to perceptual categorization, similar effects were also found in auditory regions (Figs. 4, 6). This raises the possibility that the observed effects in pitch motor areas simply reflect a downstream ‘echo’ of those in auditory regions. To scrutinize this possibility, we employed a recent mTE framework (Yu et al., 2019; Zhou et al., 2022) to measure directed information transfer between these fROIs, based on their category-related activation patterns derived from the trained classifiers (see Materials and Methods).

Figure 7 illustrates category-related information transfer between pitch motor and auditory areas (see also Table S3). Within each hemisphere, category-related information was transferred bidirectionally between the PMC and STC fROIs, along with an additional feedback connection from the PMC to AC in the right hemisphere. These findings indicate that pitch motor areas did not merely receive passive input from auditory regions, but rather actively influenced them during prosodic category processing. While the AC reciprocally exchanged information with the STC, it did not project directly to the PMC.

**Figure 7.**
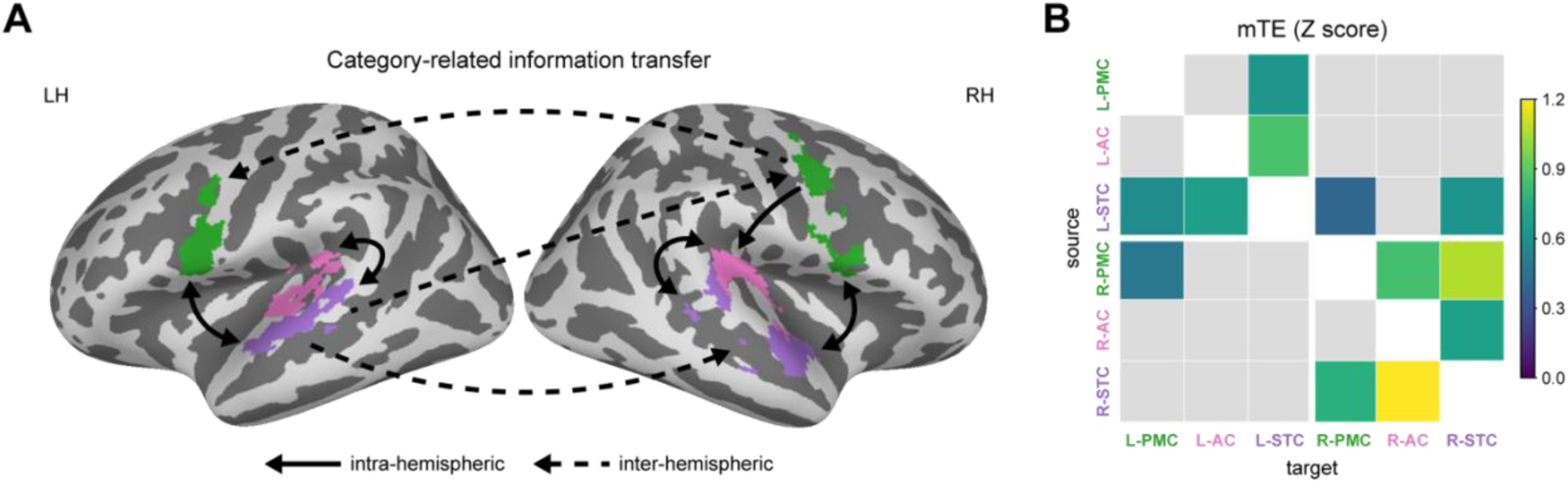
Category-related information transfer between fROIs. **A**, Directed connections showing above-chance multivariate transfer entropy (mTE) estimates (one-sample, one-sided *t*-tests against zero, FDR-corrected *q*<0.05). Solid arrows denote intra-hemispheric connections, whereas dashed arrows represent inter-hemispheric connections. **B**, mTE estimates for significant directed connections. Rows and columns show the source and target fROIs, respectively. Non-significant connections are masked in gray.

In contrast to intra-hemispheric connections, the information transfer between hemispheres was relatively sparse. Notably, the right PMC received input from the left STC (also indirectly via the right STC), and projected unidirectionally to its left homologue.

Collectively, these results demonstrate that prosodic category perception is supported by an active interplay between pitch motor and auditory areas, with right-hemispheric motor areas potentially orchestrating the exchange of category-related information across hemispheres.

## Discussion

Despite ample evidence implicating the PMC in speech processing, its functional role in perception remains debated. Here, we examined this question through prosody, a fundamental yet often overlooked aspect of speech. After localizing motor regions relevant to vocal pitch production using a simple singing task, we decoded perceived prosodic categories from neural activity patterns within these regions. The dorsal subregion, previously associated with vocal pitch modulation, contributed more strongly to decoding than the ventral subregion, particularly in the left PMC. Moreover, category decoding around stimulus offset, i.e., prior to behavioral responses, mirrored behavioral discriminability, implying the functional relation of these representations to perceptual categorization. Finally, bidirectional transfer of category-related information between pitch motor and auditory areas argues against the PMC merely ‘echoing’ processing in auditory regions. Together, these findings suggest that motor contributions to speech perception extend from articulatory control of the vocal-tract filter to laryngeal pitch control of the sound source.

### Functional localization of pitch motor areas

In contrast to prior work using functional MRI or intracranial recordings (e.g., Brown et al., 2008; Dichter et al., 2018; Eichert et al., 2020), we applied searchlight-based decoding to MEG data collected during singing on schwa (Fig. 1A), which engages laryngeal control while minimizing the movements of supralaryngeal articulators. This approach identified two PMC subregions anatomically consistent with the putative vLMC and dLMC (Fig. 3), demonstrating the feasibility of localizing motor regions relevant to vocal pitch production with MEG.

The involvement of both the putative vLMC and dLMC may reflect the two integral components of the singing task: phonation and pitch modulation. Phonation, often associated with the vLMC, generates vocal fold vibration, whereas laryngeal modulation of that vibration, more closely linked to the dLMC, produces changes in pitch (e.g., Dichter et al., 2018; Eichert et al., 2020; Hickok et al., 2023). However, the singing task may also have recruited broader vocal processes, including respiration and vocal effort (e.g., Zarate, 2013; Herbst, 2017). Therefore, the localized motor regions should be interpreted as relevant to vocal pitch production broadly rather than to pitch modulation alone. In addition, substantial inter-individual variability in PMC organization underscores the importance of subject-specific approaches (Branco et al., 2003; Gordon et al., 2023). Future localizers could use longer, subject-specific protocols and more selective contrasts to improve individual localization and better separate pitch modulation, phonation, and broader vocal processes.

### Effector-relevant motor area involvement in prosodic category perception

Decoding perceived prosodic categories from motor regions relevant to vocal pitch production is consistent with effector-relevant motor area involvement in prosody perception (Fig. 4A,D). This effect was unlikely to be explained solely by neural responses to the modeled low-level acoustic features, including stimulus envelope and pitch (Fig. 1B), although contributions from other acoustic features or higher-order auditory processing cannot be excluded. Within the localized areas, the dorsal subregion—anatomically corresponding to the putative dLMC and linked more closely to pitch modulation (e.g., Dichter et al., 2018; Finkel et al., 2019)—contributed more strongly to decoding, particularly in the left PMC (Fig. 5). Under motor theories of speech perception (Liberman and Mattingly, 1985), this dorsal contribution may reflect mapping prosodic pitch contours onto the laryngeal gestures used to produce them.

This interpretation contrasts with action prediction accounts, which attribute PMC involvement to the type of communicative response (e.g., manual or vocal) elicited by a perceived utterance (e.g., Pulvermüller et al., 2014; Tomasello, 2023). Because both statement and question utterances used here would typically elicit vocal responses, these accounts provide no clear basis for distinguishing the two prosodic categories within vocal motor regions. Even if decoding reflected a shared vocal response, the strongest contribution might be expected from the ventral subregion corresponding to the putative vLMC, more closely associated with phonation and speech sound coordination (Eichert et al., 2020; Hickok et al., 2023). Instead, the stronger dorsal contribution is more consistent with motor area involvement related to the perceived prosodic features than with the action elicited by the utterance.

The correspondence between neural and behavioral discriminability further implicates pitch motor areas in perceptual categorization (Fig. 6). Notably, this relationship was restricted to time intervals around stimulus offset, before most behavioral responses were made in the behavioral experiment, where responding was not delayed (Fig. 2B). By contrast, later decoding did not track behavioral discriminability and may instead reflect post-categorization processes, such as the maintenance of category-related representations.

Finally, bidirectional transfer of category-related information between pitch motor and auditory areas argues against PMC activity merely reflecting a downstream ‘echo’ of processing in auditory regions (Fig. 7). Together, these findings link category-related activity in pitch motor areas to the representation and categorization of prosodic percepts, in interaction with auditory regions. Nevertheless, the present MEG evidence is correlational: neither decoding, brain-behavior correlations, nor mTE establishes that pitch motor areas are *causally* necessary for prosodic category perception. Establishing their causal relevance will require lesion or neurostimulation studies.

### Neural dynamics of prosodic category processing

Our findings further revealed spatiotemporal dynamics of prosodic category processing across auditory and motor regions (Fig. 4). Perceived prosodic categories became decodable at ∼430 ms after stimulus onset, approximately 200 ms after the pitch divergence point. This time course aligns with prior findings of categorical phonemic representations, which emerge 100–200 ms after phoneme onset (e.g., Chang et al., 2010; Baek et al., 2026). This suggests that, similar to phonemes, category-related prosodic representations are formed within 200 ms after the relevant acoustic landmark, consistent with a shared processing principle underlying the neural abstraction of segmental and suprasegmental features (Baek et al., 2026).

Along with pitch motor areas, significant decoding was observed in the AC and STC, in line with previous findings of categorical prosodic representations in auditory regions (Gnanateja et al., 2025; Baek et al., 2026). Interestingly, decoding in both auditory and motor areas emerged within a similar time window around stimulus offset. These findings are consistent with parallel categorical processing across distributed cortical regions (Keshishian et al., 2023; Baek et al., 2026), rather than categorization being confined to a focal area within the posterior STC (Bouton et al., 2018). Similarly, recent evidence indicates that speech information is encoded simultaneously in frontal, including premotor, and temporal regions via parallel subcortical input (Hullett et al., 2025). Such parallel processing may support robust perception or real-time top-down modulation (e.g., Keshishian et al., 2023; Hullett et al., 2025).

Auditory regions additionally showed significant category decoding in later time windows. Because this late decoding did not mirror perceptual discriminability (Fig. 6F,G) and occurred after the time at which participants typically responded without a delay (Fig. 2B), it may reflect post-categorization processes, such as the maintenance or reinstantiation of category-related prosodic information under the delayed response requirement.

### Hemispheric asymmetry in production and perception

Although no directional lateralization was predicted or formally tested, descriptive hemispheric differences were observed in the functional localizer (Fig. 3). Singing decoding was more robust and spatially extensive in the left auditory regions than in the right, alongside a larger ventral PMC cluster in the left hemisphere. This leftward pattern may relate to left-hemispheric predominance in sensorimotor integration during vocalization (e.g., Hickok et al., 2011; Behroozmand et al., 2015). In contrast, the dorsal cluster extended more broadly in the right PMC than in the left, consistent with previous findings suggesting a greater role for the right dLMC in pitch regulation (e.g., Dichter et al., 2018; Finkel et al., 2019).

Hemispheric asymmetry during prosody perception was more nuanced. Overall, perceived prosodic categories were decodable bilaterally (Fig. 4). However, dorsal decoding importance was significantly greater than ventral importance in the left, but not right, PMC (Fig. 5B), raising the possibility of hemispheric differentiation without establishing a direct hemispheric difference. In line with this possibility, only the right PMC fROI showed task-invariant category-related representations across the prosody and phoneme tasks (Fig. S5), suggesting relatively automatic involvement and a potential fundamental scaffold for prosodic category processing (see Chevillet et al., 2013, for a similar conclusion for phonemes). Moreover, right pitch motor areas received category-related information from contralateral auditory regions and unidirectionally informed their left homologues (Fig. 7), implying that they may serve as a central hub for cross-hemispheric communication during prosodic category processing. This potential weight of the right PMC may reflect a right-hemispheric preference for intonational aspects of prosody relative to linguistic functions such as syntax and semantics (e.g., van Lancker, 1980; Gandour et al., 2004; Chien et al., 2020), or greater right-hemispheric involvement during challenging listening situations (e.g., Hartwigsen and Siebner, 2012; Liang et al., 2023). Because the hemispheres were not directly compared, future studies should determine whether these patterns reflect reliable functional asymmetries.

## Conclusion

Overall, our findings link motor regions relevant to vocal pitch production to the representation and categorization of prosodic percepts. By extending effector-relevant motor area involvement from segmental features shaped by articulators configuring the vocal-tract filter to suprasegmental features generated through the laryngeal sound source, these findings suggest a broader organizational principle for forming abstract speech representations.

## Conflict of interest

The authors declare no competing financial interests.

## Acknowledgements

This work was supported by the Max Planck Society and the Otto Hahn Award to D.S. We thank Florian Scharf for extensive efforts in stimulus generation and Yvonne Wolff-Rosier for assistance in conducting the experiments and collecting the data. We also express our gratitude to Felix Bernoully for creating the icons used in the figures.

## Author contributions

S-C.B. and D.S. designed research; S-C.B., S-G.K., B.M., and D.S. performed research; B.M. and D.S. contributed data; S-C.B. and D.S. analyzed data; S-C.B. wrote the first draft of the paper; S-C.B., S-G.K., B.M., and D.S. edited the paper; S-C.B. and D.S. wrote the paper.

## Notes

https://osf.io/4u5kx/

### Competing Interest Statement

The authors have declared no competing interest.

### Summary of Updates

Clarifying the conceptual advance and theoretical implications; Improving the methodological description and extending the analyses; Refining and tempering the interpretation

## References

Ablin P, Cardoso J-F, Gramfort A (2018) Faster independent component analysis by preconditioning with Hessian approximations. IEEE Transactions on Signal Processing 66:4040–4049.

Baek S-C, Kim S-G, Maess B, Grigutsch M, Sammler D (2026) Dynamic acoustic-to-categorical representations of phonemes and prosody along ventral and dorsal speech streams. Nat Commun 17:6082.

Behroozmand R, Shebek R, Hansen DR, Oya H, Robin DA, Howard MA, Greenlee JDW (2015) Sensory–motor networks involved in speech production and motor control: An fMRI study. Neuroimage 109:418–428.

Belyk M, Brown S (2017) The origins of the vocal brain in humans. Neurosci Biobehav Rev 77:177–193.

Boba P, Bollmann D, Schoepe D, Wester N, Wiesel J, Hamacher K (2015) Efficient computation and statistical assessment of transfer entropy. Front Phys 3:10.

Boersma P (2001) Praat, a system for doing phonetics by computer. Glot Int 5:341–345.

Bouton S, Chambon V, Tyrand R, Guggisberg AG, Seeck M, Karkar S, van de Ville D, Giraud A-L (2018) Focal versus distributed temporal cortex activity for speech sound category assignment. Proc Natl Acad Sci U S A 115:E1299–E1308.

Branco DM, Coelho TM, Branco BM, Schmidt L, Calcagnotto ME, Portuguez M, Neto EP, Paglioli E, Palmini A, Lima J V., Da Costa JC (2003) Functional variability of the human cortical motor map: Electrical stimulation findings in perirolandic epilepsy surgery. Journal of Clinical Neurophysiology 20:17–25.

Brown S, Ngan E, Liotti M (2008) A larynx area in the human motor cortex. Cerebral Cortex 18:837–845.

Chang EF, Rieger JW, Johnson K, Berger MS, Barbaro NM, Knight RT (2010) Categorical speech representation in human superior temporal gyrus. Nat Neurosci 13:1428–1432.

Chevillet MA, Jiang X, Rauschecker JP, Riesenhuber M (2013) Automatic phoneme category selectivity in the dorsal auditory stream. The Journal of Neuroscience 33:5208–5215.

Chien P, Friederici AD, Hartwigsen G, Sammler D (2020) Neural correlates of intonation and lexical tone in tonal and non-tonal language speakers. Hum Brain Mapp 41:1842–1858.

D’Ausilio A, Pulvermüller F, Salmas P, Bufalari I, Begliomini C, Fadiga L (2009) The motor somatotopy of speech perception. Current Biology 19:381–385.

de Cheveigné A (2020) ZapLine: A simple and effective method to remove power line artifacts. Neuroimage 207:116356.

DeWitt I, Rauschecker JP (2012) Phoneme and word recognition in the auditory ventral stream. Proc Natl Acad Sci U S A 109:E505–E514.

Dichter BK, Breshears JD, Leonard MK, Chang EF (2018) The control of vocal pitch in human laryngeal motor cortex. Cell 174:21–31.

Du Y, Buchsbaum BR, Grady CL, Alain C (2014) Noise differentially impacts phoneme representations in the auditory and speech motor systems. Proc Natl Acad Sci U S A 111:7126–7131.

Eichert N, Papp D, Mars RB, Watkins KE (2020) Mapping human laryngeal motor cortex during vocalization. Cerebral Cortex 30:6254–6269.

Fant Gunnar (1960) Acoustic Theory of Speech Production: With Calculations Based on X-ray Studies of Russian Articulations. Mouton.

Finkel S, Veit R, Lotze M, Friberg A, Vuust P, Soekadar S, Birbaumer N, Kleber B (2019) Intermittent theta burst stimulation over right somatosensory larynx cortex enhances vocal pitch-regulation in nonsingers. Hum Brain Mapp 40:2174–2187.

Gandour J, Tong Y, Wong D, Talavage T, Dzemidzic M, Xu Y, Li X, Lowe M (2004) Hemispheric roles in the perception of speech prosody. Neuroimage 23:344–357.

Glasser MF, Coalson TS, Robinson EC, Hacker CD, Harwell J, Yacoub E, Ugurbil K, Andersson J, Beckmann CF, Jenkinson M, Smith SM, Van Essen DC (2016) A multi-modal parcellation of human cerebral cortex. Nature 536:171–178.

Gnanateja GN, Rupp K, Llanos F, Hect J, German JS, Teichert T, Abel TJ, Chandrasekaran B (2025) Cortical processing of discrete prosodic patterns in continuous speech. Nat Commun 16:1947.

Gordon EM et al. (2023) A somato-cognitive action network alternates with effector regions in motor cortex. Nature 617:351–359.

Hartwigsen G, Siebner HR (2012) Probing the involvement of the right hemisphere in language processing with online transcranial magnetic stimulation in healthy volunteers. Aphasiology 26:1131–1152.

Haufe S, Meinecke F, Görgen K, Dähne S, Haynes J-D, Blankertz B, Bießmann F (2014) On the interpretation of weight vectors of linear models in multivariate neuroimaging. Neuroimage 87:96–110.

Hautus MJ (1995) Corrections for extreme proportions and their biasing effects on estimated values of d′. Behavior Research Methods, Instruments, & Computers 27:46–51.

Haynes J-D, Rees G (2006) Decoding mental states from brain activity in humans. Nat Rev Neurosci 7:523–534.

Hellbernd N, Sammler D (2016) Prosody conveys speaker’s intentions: Acoustic cues for speech act perception. J Mem Lang 88:70–86.

Herbst CT (2017) A review of singing voice subsystem interactions—toward an extended physiological model of “support.” Journal of Voice 31:249.e13–249.e19.

Hickok G (2010) The role of mirror neurons in speech perception and action word semantics. Lang Cogn Process 25:749–776.

Hickok G, Houde J, Rong F (2011) Sensorimotor integration in speech processing: Computational basis and neural organization. Neuron 69:407–422.

Hickok G, Poeppel D (2007) The cortical organization of speech processing. Nat Rev Neurosci 8:393–402.

Hickok G, Venezia J, Teghipco A (2023) Beyond Broca: neural architecture and evolution of a dual motor speech coordination system. Brain 146:1775–1790.

Hullett PW, Leonard MK, Gorno-Tempini ML, Mandelli ML, Chang EF (2025) Parallel encoding of speech in human frontal and temporal lobes. Nat Commun 17:814.

Jas M, Engemann DA, Bekhti Y, Raimondo F, Gramfort A (2017) Autoreject: Automated artifact rejection for MEG and EEG data. Neuroimage 159:417–429.

Kaernbach C (1991) Simple adaptive testing with the weighted up-down method. Percept Psychophys 49:227–229.

Kawahara H, Mutsui H (2003) Auditory morphing based on an elastic perceptual distance metric in an interference-free time-frequency representation. In: IEEE International Conference on Acoustics, Speech, and Signal Processing, 2003. Proceedings.(ICASSP’03), pp I–I. Hong Kong: IEEE.

Keshishian M, Akkol S, Herrero J, Bickel S, Mehta AD, Mesgarani N (2023) Joint, distributed and hierarchically organized encoding of linguistic features in the human auditory cortex. Nat Hum Behav 7:740–753.

Kingdom FAA, Prins N (2016) Psychophysics. Elsevier.

Kriegeskorte N, Goebel R, Bandettini P (2006) Information-based functional brain mapping. Proc Natl Acad Sci U S A 103:3863–3868.

Kurumada C, Roettger TB (2022) Thinking probabilistically in the study of intonational speech prosody. Wiley Interdiscip Rev Cogn Sci 13:e1579.

Ledoit O, Wolf M (2004) A well-conditioned estimator for large-dimensional covariance matrices. J Multivar Anal 88:365–411.

Liang B, Li Y, Zhao W, Du Y (2023) Bilateral human laryngeal motor cortex in perceptual decision of lexical tone and voicing of consonant. Nat Commun 14:4710.

Liberman AM, Mattingly IG (1985) The motor theory of speech perception revised. Cognition 21:1–36.

Lu J, Li Y, Zhao Z, Liu Y, Zhu Y, Mao Y, Wu J, Chang EF (2023) Neural control of lexical tone production in human laryngeal motor cortex. Nat Commun 14:6917.

Maess B, Schröger E, Widmann A (2016) High-pass filters and baseline correction in M/EEG analysis-continued discussion. J Neurosci Methods 266:171–172.

Maris E, Oostenveld R (2007) Nonparametric statistical testing of EEG- and MEG-data. J Neurosci Methods 164:177–190.

Möttönen R, Dutton R, Watkins KE (2013) Auditory-motor processing of speech sounds. Cerebral Cortex 23:1190–1197.

Möttönen R, Watkins KE (2009) Motor representations of articulators contribute to categorical perception of speech sounds. The Journal of Neuroscience 29:9819–9825.

Nonyane BAS, Theobald CM (2007) Design sequences for sensory studies: Achieving balance for carry-over and position effects. British Journal of Mathematical and Statistical Psychology 60:339–349.

Oldfield RC (1971) The assessment and analysis of handedness: The Edinburgh inventory. Neuropsychologia 9:97–113.

Perron M, Vuong V, Grassi MW, Imran A, Alain C (2024) Engagement of the speech motor system in challenging speech perception: Activation likelihood estimation meta-analyses. Hum Brain Mapp 45:e70023.

Pierrehumbert J, Hirschberg J (1990) The meaning of intonational contours in the interpretation of discourse. In: Intentions in Communication (Cohen PR, Morgan J, Pollack ME, eds). Cambridge, MA: The MIT Press.

Pulvermüller F, Moseley RL, Egorova N, Shebani Z, Boulenger V (2014) Motor cognition–motor semantics: Action perception theory of cognition and communication. Neuropsychologia 55:71–84.

Sammler D, Grosbras MH, Anwander A, Bestelmeyer PEG, Belin P (2015) Dorsal and ventral pathways for prosody. Current Biology 25:3079–3085.

Sekihara K, Nagarajan SS (2008) Adaptive spatial filters. In: Adaptive Spatial Filters for Electromagnetic Brain Imaging, pp 37–63. Berlin Heidelberg: Springer.

Si X, Zhou W, Hong B (2017) Cooperative cortical network for categorical processing of Chinese lexical tone. Proc Natl Acad Sci U S A 114:12303–12308.

Takens F (1981) Detecting strange attractors in turbulence. In: Dynamical Systems and Turbulence, Warwick 1980 (Rand DA, Young L-S, eds), pp 366–381. Springer, Berlin, Heidelberg.

Tomasello R (2023) Linguistic signs in action: The neuropragmatics of speech acts. Brain Lang 236:105203.

Tomasello R, Grisoni L, Boux I, Sammler D, Pulvermüller F (2022) Instantaneous neural processing of communicative functions conveyed by speech prosody. Cerebral Cortex 32:4885–4901.

van Lancker D (1980) Cerebral lateralization of pitch cues in the linguistic signal. Paper in Linguistics 13:201–277.

Van Veen BD, Van Drongelen W, Yuchtman M, Suzuki A (1997) Localization of brain electrical activity via linearly constrained minimum variance spatial filtering. IEEE Trans Biomed Eng 44:867–880.

Wallot S, Mønster D (2018) Calculation of average mutual information (AMI) and false-nearest neighbors (FNN) for the estimation of embedding parameters of multidimensional time series in Matlab. Front Psychol 9:1679.

Westner BU, Dalal SS, Gramfort A, Litvak V, Mosher JC, Oostenveld R, Schoffelen J-M (2022) A unified view on beamformers for M/EEG source reconstruction. Neuroimage 246:118789.

Wibral M, Rahm B, Rieder M, Lindner M, Vicente R, Kaiser J (2011) Transfer entropy in magnetoencephalographic data: Quantifying information flow in cortical and cerebellar networks. Prog Biophys Mol Biol 105:80–97.

Wichmann FA, Hill NJ (2001) The psychometric function: I. Fitting, sampling, and goodness of fit. Percept Psychophys 63:1293–1313.

Widmann A, Schröger E, Maess B (2015) Digital filter design for electrophysiological data – a practical approach. J Neurosci Methods 250:34–46.

Xie X, Buxó-Lugo A, Kurumada C (2021) Encoding and decoding of meaning through structured variability in intonational speech prosody. Cognition 211:104619.

Yu S, Sanchez Giraldo LG, Jenssen R, Principe JC (2019) Multivariate extension of matrix-based Renyi’s α-order entropy functional. IEEE Trans Pattern Anal Mach Intell 42:1–1.

Zarate JM (2013) The neural control of singing. Front Hum Neurosci 7:48369.

Zhou W, Yu S, Chen B (2022) Causality detection with matrix-based transfer entropy. Inf Sci 613:357–375.

