## Supplemental Material for "Pitch motor areas contribute to the perception of prosodic categories in speech"

### Supplemental materials

### Supplemental Figures

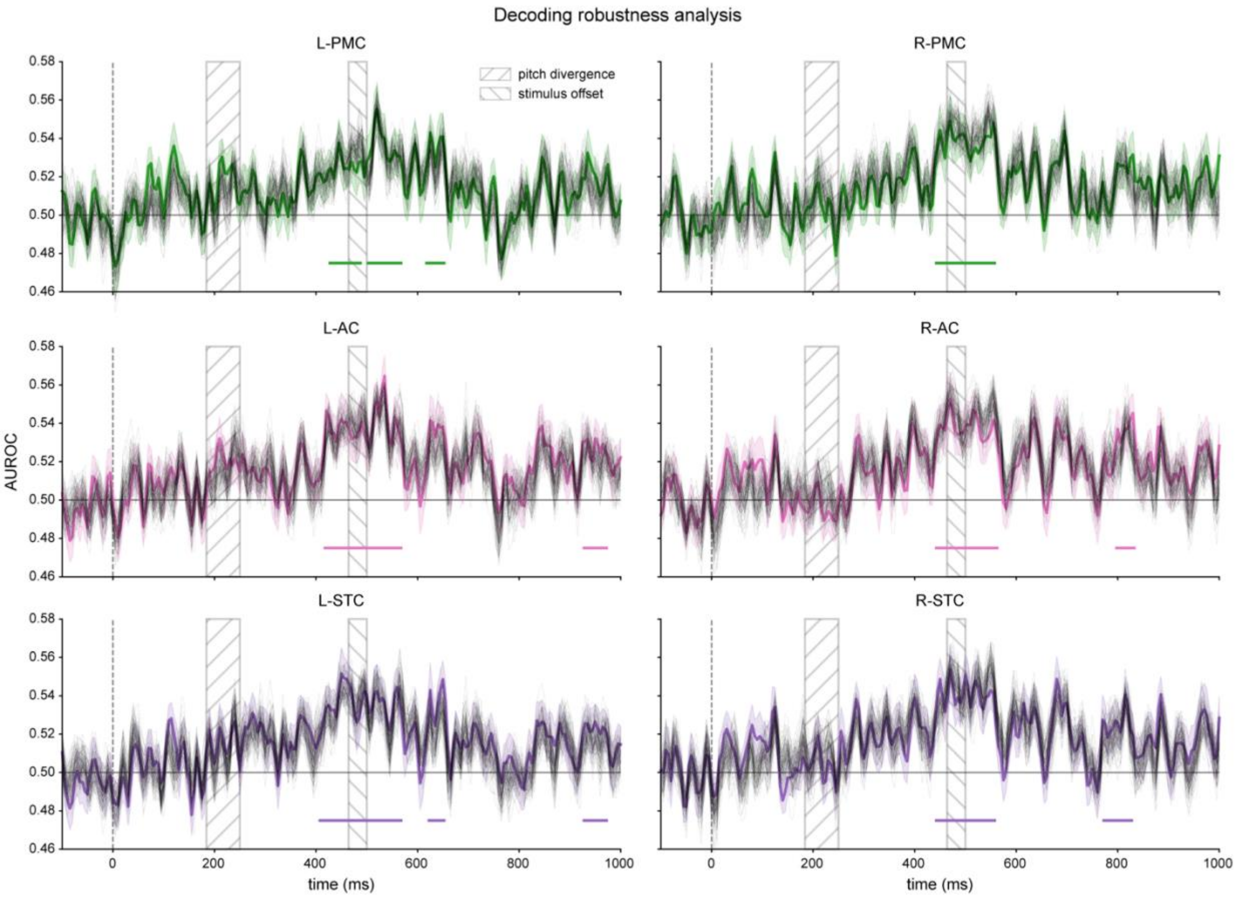

**Figure S1.** Decoding robustness analysis. To determine whether the effects shown in Figure 4 depended on a particular probabilistically generated set of prosody pseudo-labels, we regenerated the pseudo-labels using 100 different random seeds (1-100) and repeated the same decoding analysis for each realization. Decoding models were trained to decode prosody on all trials without behavioral responses and tested on trials with overt responses from the prosody task. Left- (L-) and right-hemispheric (R-) functional regions of interest (fROIs) are shown in the left and right columns, respectively. Time-resolved decoding performance is quantified as the area under the receiver operating characteristic curve (AUROC). Thin black lines show the group-averaged decoding time series for the 100 pseudo-label realizations. The original decoding results reported in Figure 4 are replotted as colored lines, with shaded areas indicating the standard error of the mean (SEM). Horizontal lines above the x-axis denote the significant time intervals identified in Figure 4. The rectangular areas with rising hatching (left) and falling hatching (right) represent the ranges of the pitch divergence point and stimulus offset, respectively. Across all fROIs, the time series obtained with different pseudo-label realizations largely fell within the SEM of the original results, indicating that decoding performance was robust to the stochastic generation of pseudo-labels. PMC, premotor cortex; AC, auditory cortex; STC, superior temporal cortex.

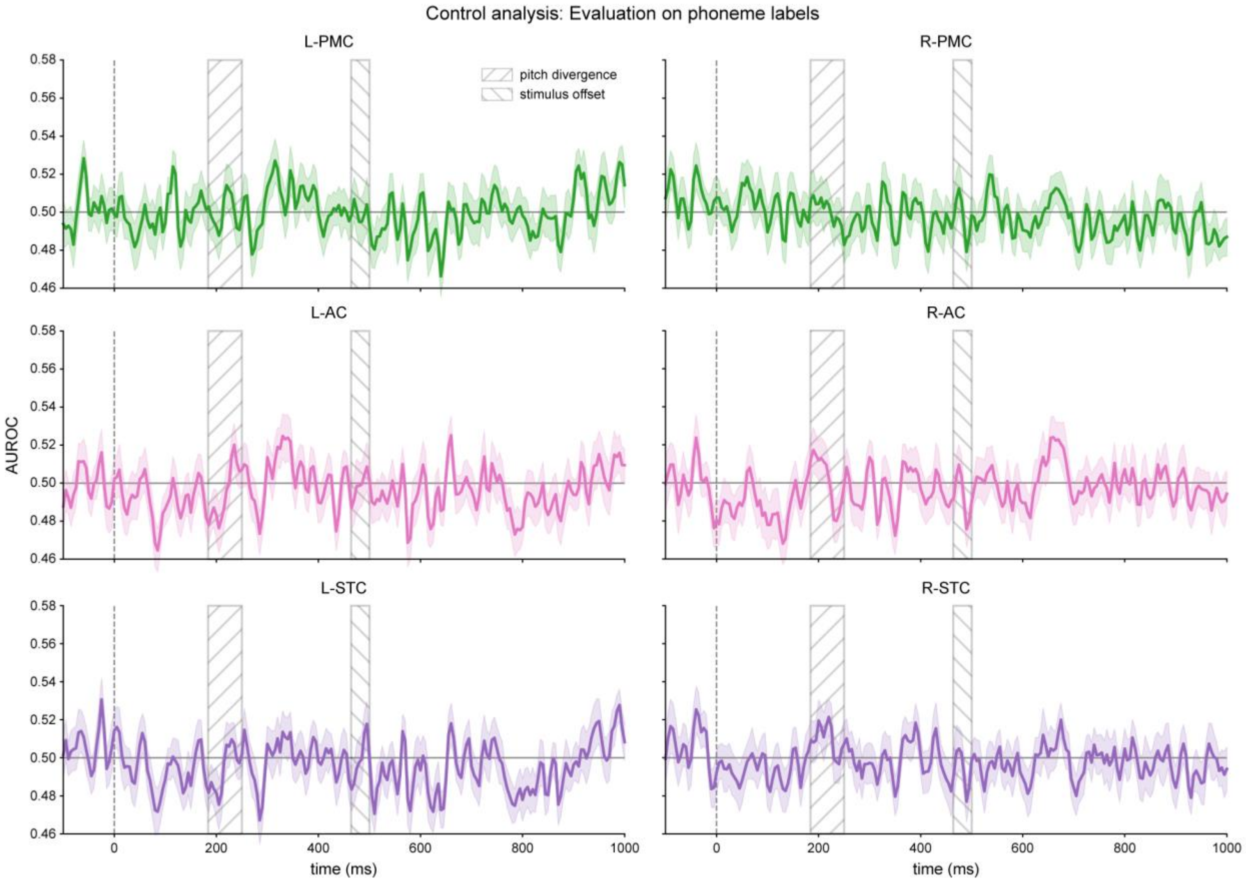

**Figure S2.** Control analysis: Evaluation on phoneme labels. To test whether the effects in Figure 4 were specific to prosody, the same decoding models—trained to decode prosody on all trials without behavioral responses—were tested on trials with overt responses from the phoneme task. Time-resolved decoding performance, quantified as AUROC, is shown for the left- (left column) and right-hemispheric (right column) fROIs. Shaded areas indicate SEM. No fROIs showed significant decoding of phonemic categories (one-sample, one-sided $t$-tests against 0.5, cluster-forming threshold (CFT) at $p < 0.05$, family-wise error rate (FWER) controlled at 0.05 across time points per fROI; all cluster $ps > 0.05$). All other plotting details are the same as Figure S1.

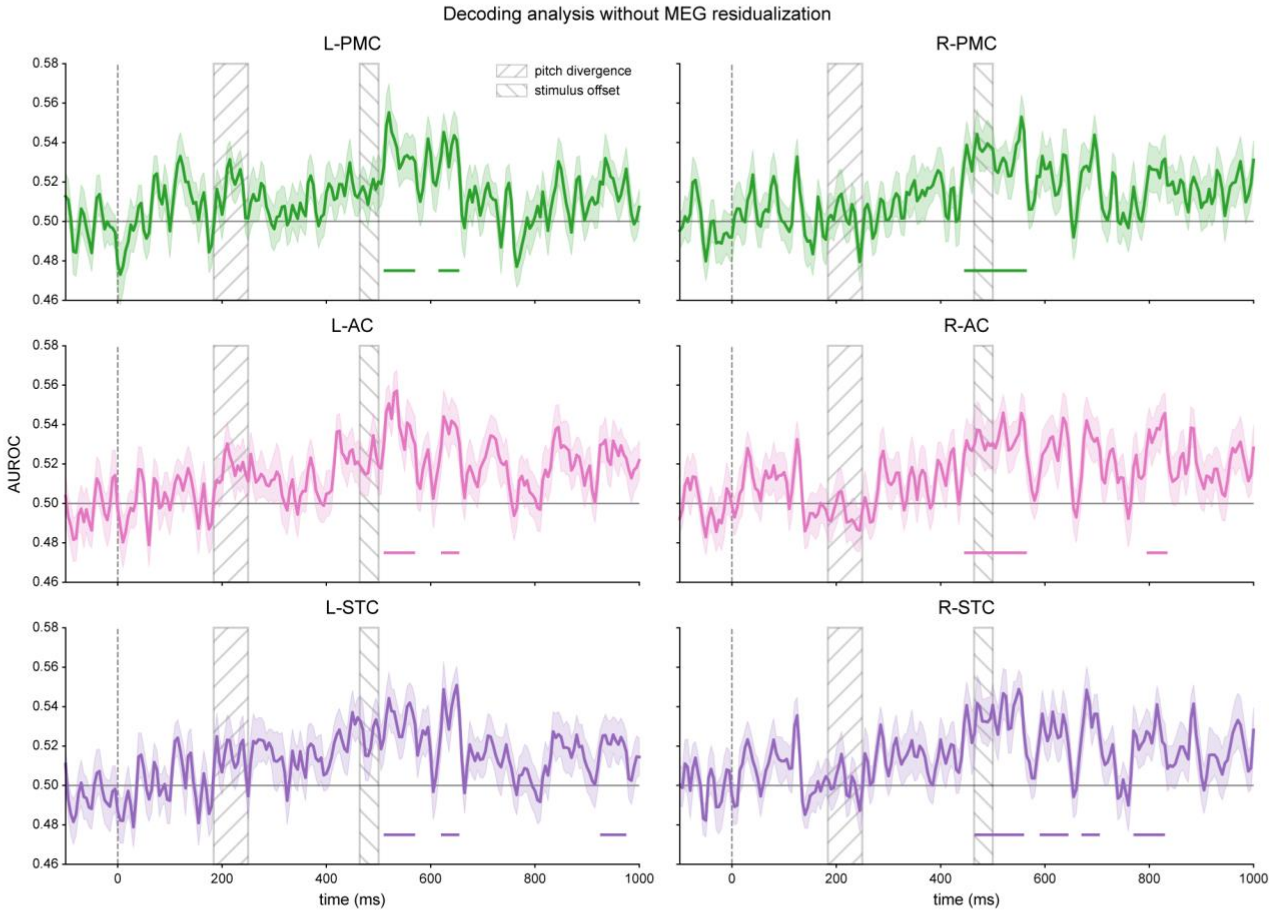

**Figure S3.** Decoding analysis without MEG residualization. To assess whether the effects shown in Figure 4 were primarily explained by low-level acoustic processing, we repeated the same decoding analysis using MEG source signals without regressing out neural responses to the modeled acoustic features (see Fig. 1B). As in Figure 4, decoding models were trained on all trials without behavioral responses and then tested on trials with overt responses from the prosody task. Horizontal lines above the x-axis denote significant time intervals (statistical inference as in Fig. 4). All other plotting details are the same as Figure S1. Compared to Figure 4, decoding performance was reduced and significant for a shorter duration before and around stimulus offset, This suggests that responses to the modeled low-level acoustic features may partially obscure neural patterns associated with perceived prosodic categories, arguing against a purely acoustic account of the effects observed in Figure 4.

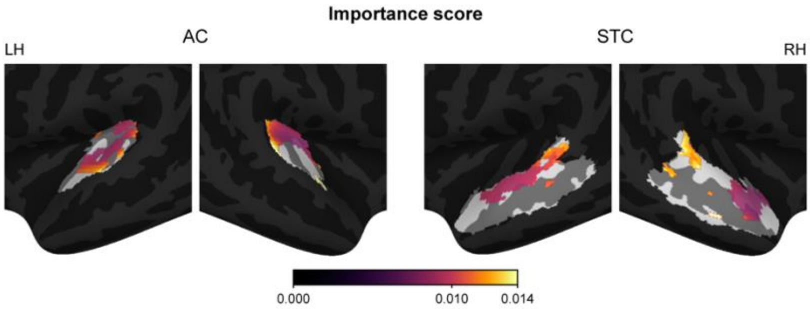

**Figure S4.** Group-averaged vertex-wise importance scores for decoding within the AC and STC in the left (LH) and right (RH) hemispheres.

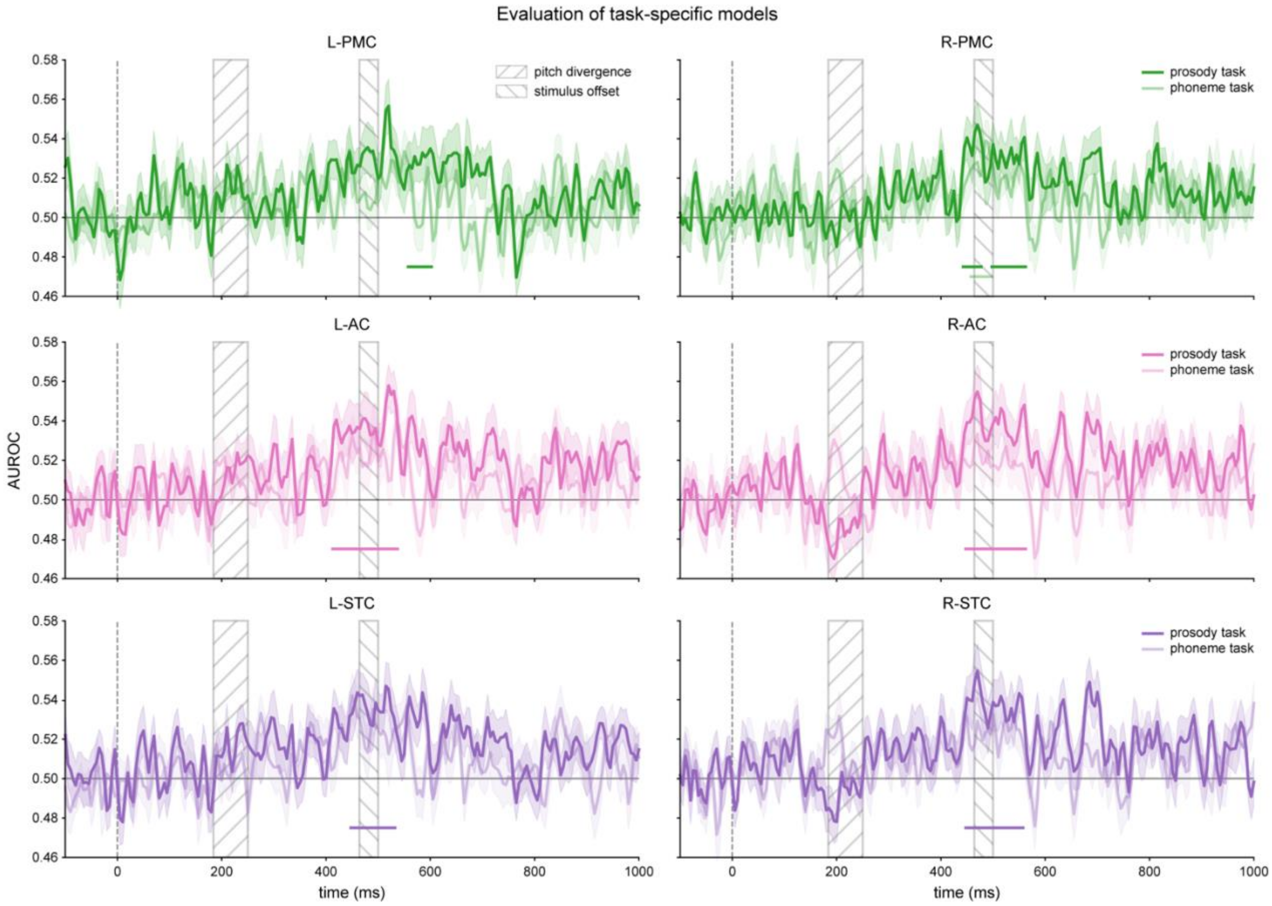

**Figure S5.** Task-specific model evaluation. In the follow-up decoding analyses, decoding models were trained on separate subsets of trials without behavioral responses from either the prosody (vivid lines) or the phoneme (pale lines) identification task, using prosody pseudo-labels. Both task-specific models were then tested on trials with overt responses from the prosody task. Time-resolved decoding performance, quantified as AUROC, is shown for the left- (left column) and right-hemispheric (right column) fROIs. Shaded areas indicate SEM. Horizontal lines above the x-axis denote significant time intervals (one-sample, one-sided $t$-tests against 0.5, CFT at $p < 0.05$, FWER rate controlled at 0.05 across time points per fROI, fROI-wise Bonferroni-corrected $p < 0.0042$). All other plotting details are the same as Figure S1. Direct comparisons between the two task-specific models did not yield significant modulation effects (paired, one-sided $t$-tests against zero, CFT at $p < 0.05$, FWER controlled at 0.05 across time points per fROI; all cluster $ps > 0.05$).

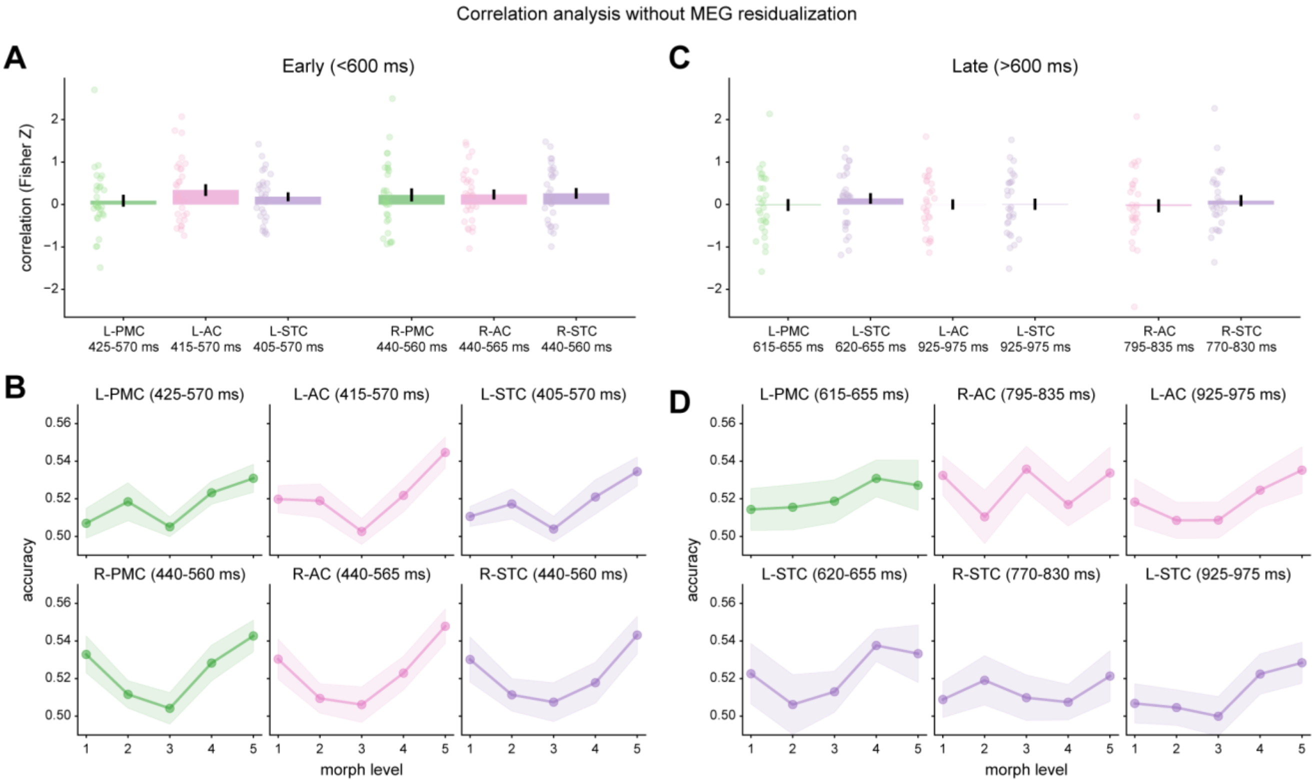

**Figure S6.** Correlation analysis without MEG residualization. The same analysis as in Figure 6D-G was performed on the MEG source signals without regressing out responses to the modeled low-level acoustic features (see Fig. 1B), using the identical time intervals. Statistical inference and plotting details are the same as in Figure 6D-G. **A,C**, Correlation (Fisher-Z transformed) between level-wise decoding accuracy and behavioral discriminability across fROIs within the time intervals before (**A**) and after (**C**) 600 ms relative to stimulus onset. No significant correlations were found in any interval (all false-discovery-rate-corrected $qs > 0.05$). **B,D**, Level-wise decoding accuracy within the time intervals before (**B**) and after (**D**) 600 ms. While decoding profiles were largely unchanged in the intervals after 600 ms, the earlier intervals showed no clear or only a weaker U-shaped pattern. These results suggest that the effects observed in Figure 6D,E are unlikely to be explained simply by neural responses to the modeled acoustic features.

### Supplemental Tables

**Table S1.** Summary statistics of the stimuli used in the MEG experiment.

| Speaker | | Morph step | Duration (ms) | Fundamental frequency ($F_0$) | | | Intensity | |
|---|---|---|---|---|---|---|---|---|
| | | | | Mean (Hz) | Range (Hz) | Slope (st/s) | Mean (dB) | Range (dB) |
| Female | S | 24 ± 6.4 | 479 ± 6.3 | 173 ± 6 | 38 ± 16 | 22 ± 4 | 65 ± 0.4 | 22 ± 1.6 |
| | ↕ | 28 ± 7.5 | 479 ± 6.3 | 177 ± 8 | 49 ± 25 | 25 ± 6 | 66 ± 0.4 | 21 ± 1.5 |
| | | 32 ± 8.7 | 479 ± 6.3 | 182 ± 10 | 64 ± 36 | 28 ± 8 | 66 ± 0.4 | 21 ± 1.8 |
| | | 36 ± 10.1 | 479 ± 6.3 | 187 ± 13 | 82 ± 48 | 33 ± 11 | 66 ± 0.4 | 21 ± 2.0 |
| | Q | 40 ± 11.6 | 479 ± 6.3 | 192 ± 16 | 102 ± 61 | 37 ± 13 | 66 ± 0.4 | 20 ± 2.1 |
| Male | S | 23 ± 9.1 | 476 ± 6.0 | 101 ± 5 | 27 ± 9 | 20 ± 5 | 64 ± 0.5 | 25 ± 1.6 |
| | ↕ | 28 ± 10.1 | 476 ± 6.0 | 104 ± 5 | 34 ± 14 | 23 ± 6 | 64 ± 0.6 | 25 ± 1.6 |
| | | 33 ± 11.2 | 476 ± 6.0 | 107 ± 6 | 43 ± 21 | 26 ± 8 | 65 ± 0.7 | 24 ± 1.8 |
| | | 37 ± 12.4 | 476 ± 6.0 | 110 ± 8 | 53 ± 27 | 30 ± 10 | 65 ± 0.7 | 24 ± 2.2 |
| | Q | 42 ± 13.8 | 476 ± 6.0 | 113 ± 9 | 65 ± 34 | 34 ± 12 | 65 ± 0.7 | 23 ± 2.4 |

Summary statistics (mean ± SD) of the acoustic properties for stimuli across the five prosody levels are reported for each speaker. Morph step indicates the stimulus position on the original 61-step prosody continuum. st/s, semitones per second; S, statement; Q, question.

**Table S2.** Cluster statistics of task-specific models for significant decoding of perceived prosodic categories.

| **Training task** | **fROI** | **Duration (ms)** | **Bonferroni $p$-value** | **Average decoding performance (AUROC ± SEM)** | **Average $t_{28}$-value** |
|---|---|---|---|---|---|
| Prosody | L-PMC | 555-605 | 0.048 | 0.529 ± 0.007 | 2.42 |
| | L-AC | 410-540 | 0.001 | 0.537 ± 0.006 | 3.29 |
| | L-STC | 445-535 | 0.001 | 0.536 ± 0.007 | 2.97 |
| | R-PMC | 440-480 | 0.023 | 0.537 ± 0.007 | 3.26 |
| | | 495-565 | 0.002 | 0.531 ± 0.006 | 2.68 |
| | R-AC | 445-565 | 0.001 | 0.539 ± 0.006 | 3.24 |
| | R-STC | 445-560 | 0.001 | 0.537 ± 0.006 | 3.09 |
| Phoneme | R-PMC | 455-500 | 0.023 | 0.528 ± 0.006 | 2.54 |

Statistics for significant temporal clusters are reported for each functional region of interest (fROI), when classifiers were trained separately on the prosody and phoneme task subsets. Average decoding performance and average $t$-values represent the grand mean across time points within each significant cluster. L-, left; R-, right; PMC, premotor cortex; AC, auditory cortex; STC, superior temporal cortex; AUROC, area under the receiver operating characteristic curve; SEM, standard error of the mean.

**Table S3.** Statistics for significant multivariate transfer entropy (mTE) estimates between functional regions of interest (fROIs).

| Source fROI | Target fROI | mTE value ± SEM | $t_{28}$-value | FDR $q$-value |
|---|---|---|---|---|
| L-PMC | L-STC | 0.631 ± 0.205 | 3.03 | 0.016 |
| L-AC | L-STC | 0.857 ± 0.254 | 3.32 | 0.010 |
| L-STC | L-PMC | 0.580 ± 0.209 | 2.73 | 0.018 |
| | L-AC | 0.668 ± 0.272 | 2.41 | 0.028 |
| | R-PMC | 0.398 ± 0.157 | 2.49 | 0.026 |
| | R-STC | 0.619 ± 0.212 | 2.87 | 0.018 |
| R-PMC | L-PMC | 0.480 ± 0.170 | 2.78 | 0.018 |
| | R-AC | 0.844 ± 0.315 | 2.63 | 0.020 |
| | R-STC | 1.062 ± 0.309 | 3.38 | 0.010 |
| R-AC | R-STC | 0.682 ± 0.247 | 2.72 | 0.018 |
| R-STC | R-PMC | 0.761 ± 0.198 | 3.77 | 0.006 |
| | R-AC | 1.308 ± 0.229 | 5.62 | <0.001 |

Transfer of category-related information from source to target within each pair of fROIs was measured using mTE. Statistics are reported for all connections that reached significance. L-, left; R-, right; PMC, premotor cortex; AC, auditory cortex; STC, superior temporal cortex; SEM, standard error of the mean; FDR, false discovery rate.
